# Learning antibody sequence constraints from allelic inclusion

**DOI:** 10.1101/2024.10.22.619760

**Authors:** Milind Jagota, Chloe Hsu, Thomas Mazumder, Kevin Sung, William S. DeWitt, Jennifer Listgarten, Frederick A. Matsen, Chun Jimmie Ye, Yun S. Song

**Affiliations:** Computer Science Division, UC Berkeley, Berkeley, CA USA; Division of Rheumatology, Department of Medicine, UCSF, San Francisco, CA, USA; Public Health Sciences Division, Fred Hutchinson Cancer Research Center, Seattle, Washington, USA; Chan Zuckerberg Biohub, San Francisco, CA, USA; Parker Institute for Cancer Immunotherapy, UCSF, San Francisco, CA, USA; Institute for Human Genetics, UCSF, San Francisco, CA, USA; Bakar Computational Health Sciences Institute, UCSF, San Francisco, California, USA; Department of Epidemiology and Biostatistics, UCSF, San Francisco, CA, USA; Department of Statistics, UC Berkeley, Berkeley, CA, USA October 23, 2024

## Abstract

Antibodies and B-cell receptors (BCRs) are produced by B cells, and are built of a heavy chain and a light chain. Although each B cell could express two different heavy chains and four different light chains, usually only a unique pair of heavy chain and light chain is expressed—a phenomenon known as *allelic exclusion*. However, a small fraction of naive-B cells violate allelic exclusion by expressing two productive light chains, one of which has impaired function; this has been called *allelic inclusion*. We demonstrate that these B cells can be used to learn constraints on antibody sequence. Using large-scale single-cell sequencing data from humans, we find examples of light chain allelic inclusion in thousands of naive-B cells, which is an order of magnitude larger than existing datasets. We train machine learning models to identify the abnormal sequences in these cells. The resulting models correlate with antibody properties that they were not trained on, including polyreactivity, surface expression, and mutation usage in affinity maturation. These correlations are larger than what is achieved by existing antibody modeling approaches, indicating that allelic inclusion data contains useful new information. We also investigate the impact of similar selection forces on the heavy chain in mouse, and observe that pairing with the surrogate light chain significantly restricts heavy chain diversity.

## Introduction

B cells are a crucial component of the adaptive immune system and they carry out many of their characteristic functions via the B-cell receptor (BCR) [1]. The BCR recognizes antigens through a membrane bound immunoglobulin (Ig) molecule, made up of two antigen-binding fragments, each containing a heavy (H) chain and a light (L) chain. Some B cells can secrete Ig molecules as antibodies in response to the presence of antigens. In humans, although each B cell could genetically express two different H chains and four different L chains (two *κ* and two *λ*), usually only a unique pair of H chain and L chain is expressed in each B cell, known as the *allelic exclusion* phenomenon (Figure 1) [2]. Antibodies are also of interest as experimental tools and therapeutics, because they can be engineered to bind desired targets [3].

**Figure 1:**
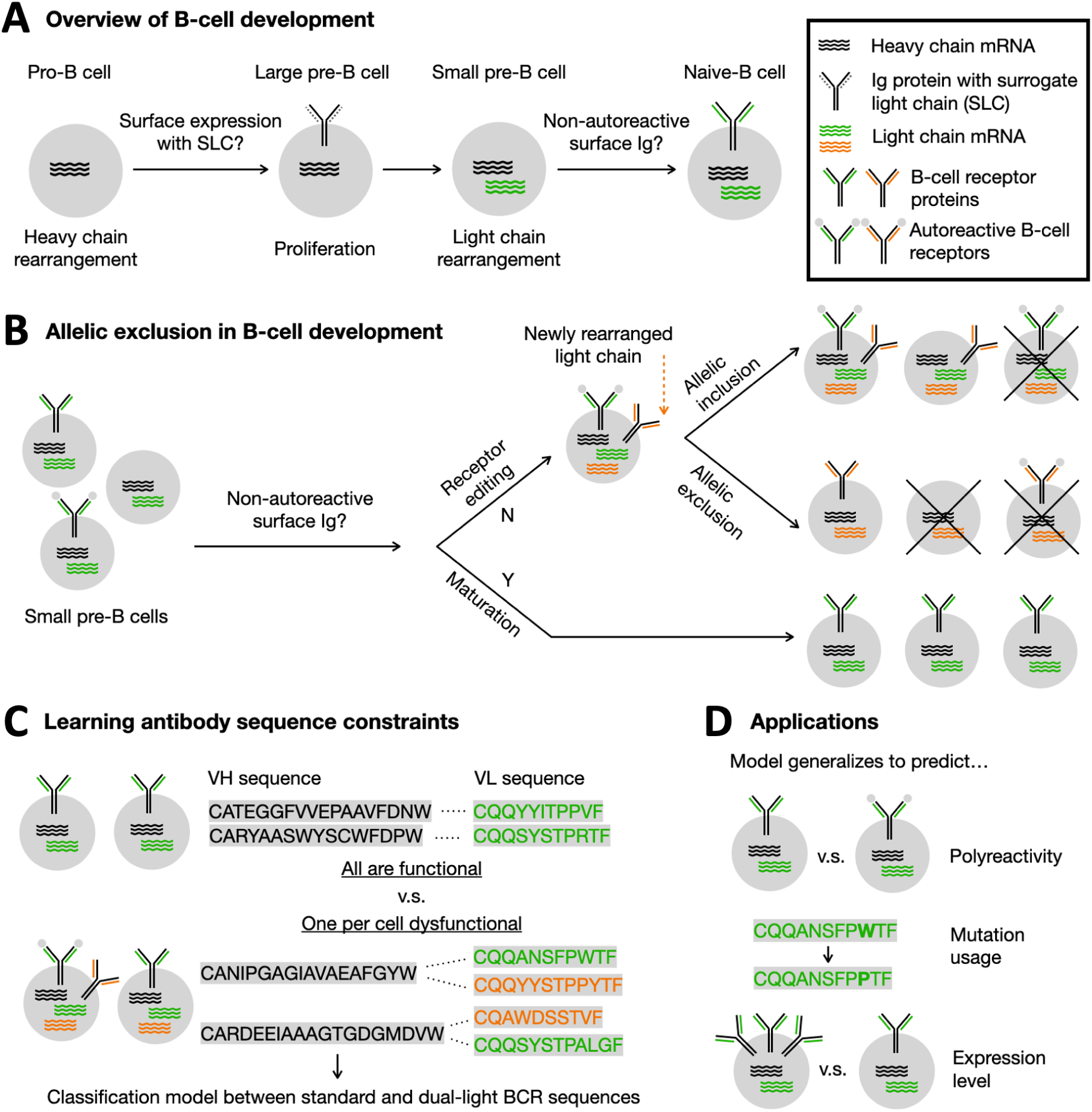
Learning antibody sequence constraints from allelic inclusion. **(A)** Overview of B-cell development. Random heavy and light chains are generated separately, with each being tested for functionality in sequential checkpoints. The heavy chain is generated in pro-B cells and must express with the surrogate light chain (SLC). The light chain is generated in pre-B cells and must express with the heavy chain with low autoreactivity. **(B)** Overview of light-chain allelic exclusion in B-cell development. Light chains are initially generated in small pre-B cells. If the initial light chain does not express on the surface or is autoreactive, new rearrangements are attempted via receptor editing. Usually only a final, successful pair of heavy and light chains is expressed as mRNA (allelic exclusion) but sometimes two attempts can be observed, one of which is autoreactive or not expressed on the surface (allelic inclusion). Cells that are still missing non-autoreactive surface BCR after receptor editing will die. **(C)** We develop machine learning models that classify antibody sequences as coming from an allelically included B cell or not. **(D)** Machine learning models of allelic inclusion generalize to predict antibody properties they were not trained on, including polyreactivity, mutation usage, and surface expression level.

An important problem in immunology and antibody engineering is to describe the constraints on antibody sequences imposed by properties like stability and polyreactivity. Although the theoretical space of antibodies is massive, most sequences are either unstable or highly polyreactive [2, 4]. These antibodies are generally not useful and can be associated with disease [5]. Our understanding of these properties has been limited by a lack of *negative examples* of sequences that violate such constraints, which we hereafter refer to as *dysfunctional*. The natural immune system filters dysfunctional BCRs via sequential checkpoints in B-cell development, occurring primarily in the bone marrow (Figure 1A). It is possible to sequence the distribution of BCRs before and after these checkpoints to observe what kinds of BCRs satisfy constraints [4, 6]. However, the throughput of such experiments in humans has been very low because of the difficulty in collecting bone marrow samples. There has also been work on measuring stability and polyreactivity via *in vitro* assays, but such assays are typically much lower throughput than binding assays and may differ from *in vivo* constraints [7–10].

In this work, we propose a new source of dysfunctional BCRs to use as negative examples for modeling antibody constraints. To observe dysfunctional sequences, we use B cells that violate the allelic exclusion phenomenon. During B-cell development, B cells first generate a heavy (H) and then a light (L) chain for the BCR (Figure 1A). These sequences must properly express, pair, and display limited autoreactivity. At each stage, if a rearrangement fails, additional rearrangements can be attempted in a process known as *receptor editing* [11]. The allelic exclusion phenomenon specifies that most B cells only express the final, successful rearrangements of the H and L chains. However, a small fraction (∼1%) of naive-B cells in humans have been shown to express two productive L chains at the level of mRNA [2], a phenomenon referred to as *allelic inclusion* (Figure 1B). When allelic inclusion occurs, one of the two transcripts has low surface expression or is autoreactive [12, 13]. Notably, naive-B cells circulate in peripheral blood, and can therefore be sequenced at much higher throughput than early B-cell precursors in the bone marrow [14–17]. Allelic inclusion has also been reported in memory-B cells, but this is much less well-characterized [18].

We sought to use dysfunctional rearrangements from allelically included B cells as negative training examples for a machine learning model (Figure 1C). Previous work observing allelic inclusion has only characterized it at a small scale (hundreds of B cells) [12, 13, 18, 19]. By leveraging recent, large single-cell BCR repertoires in humans, we were able to observe light-chain allelic inclusion at a much larger scale (several thousand B cells). However, we do not know *a priori* which sequence in each cell is dysfunctional. We developed a framework to train models on allelic inclusion data without this information and can then use the trained models to score individual sequences. The model scores generalize to predict several antibody properties associated with dysfunctionality with no additional training. These properties include polyreactivity, surface expression, and mutation usage in affinity maturation (Figure 1D).

Previous work has attempted to understand constraints on antibody sequence by only modeling functional BCR sequences that passed developmental checkpoints [14, 20–22]. However, the distribution of such sequences is shaped by both the V(D)J recombination process and the development checkpoints. A BCR sequence could appear unlikely in the distribution of functional BCRs either because it is unlikely to be generated by V(D)J recombination or because it is dysfunctional. Models that only examine functional sequences and no negative examples cannot distinguish these scenarios. Another line of work has proposed simulating the V(D)J recombination process *in silico* to generate BCR repertoires that are not subject to any selection pressures [23, 24]. This work uses unproductive BCR sequences from naive-B cells (those that are out-of-frame or have a stop codon) to fit the parameters of the simulation. However, the accuracy of this approach has not been assessed on a pre-selection repertoire of productive BCR sequences, because a sufficiently large repertoire of this kind has not previously existed. We compare our new models against these approaches and show improved ability to predict light chain sequence constraints, demonstrating the value of learning from dysfunctional examples from allelic inclusion.

Our models trained on light chain allelic inclusion are not applicable for heavy chain constraints, but we explore the impact of similar selection forces on the heavy chain using bulk repertoires in mice. We observed evidence that surface expression significantly constrains heavy-chain diversity through pairing with the surrogate light chain.

## Results

### Finding dysfunctional BCRs by observing allelic inclusion at scale

We looked to use dysfunctional BCRs observed in allelic inclusion as training data for a machine learning model. However, existing evidence of allelic inclusion in naive-B cells were based on low-throughput sequencing, with each analysis involving hundreds of B cells at most [12]. To obtain a larger training dataset, we examined recent single-cell BCR repertoires that include hundreds of thousands of B cells. In particular, a recent collection of single-cell repertoires from Jaffe *et al.* includes over 400,000 human naive-B cells [15], while another collection from van der Wijst *et al.* includes over 30,000 human naive-B cells [25]. A technical challenge for analyzing these data is the existence of *multiplet* errors. Multiplet errors occur during single-cell sequencing when two B cells are erroneously labeled with the same barcode by falling into the same droplet. A multiplet that includes two standard, allelically excluded B cells could manifest as a single barcode being associated with multiple BCR transcripts. The smaller dataset from van der Wijst *et al.* includes single-cell gene expression vectors, which were used to identify and filter multiplets computationally (**Methods**). However, the larger dataset from Jaffe *et al.* does not include gene expression vectors and has not had multiplets filtered out. The van der Wijst *et al.* dataset therefore gives a more accurate estimate of allelic inclusion frequencies.

Table 1 shows the fraction of barcodes with different counts of heavy and light chain BCR transcripts. In both datasets, the vast majority of barcodes exhibit a unique pair of heavy and light chain transcripts - as expected for the standard allelic exclusion behavior. Both datasets have about 5% of barcodes associated with two light chains and one heavy chain (referred to as *double-light*). The filtered dataset (Table 1, right) has very few barcodes associated with two heavy chains, while the unfiltered dataset (Table 1, left) has nearly 8% of barcodes associated with two heavy chains, almost equally split between having one or two light chains.

**Table 1:**
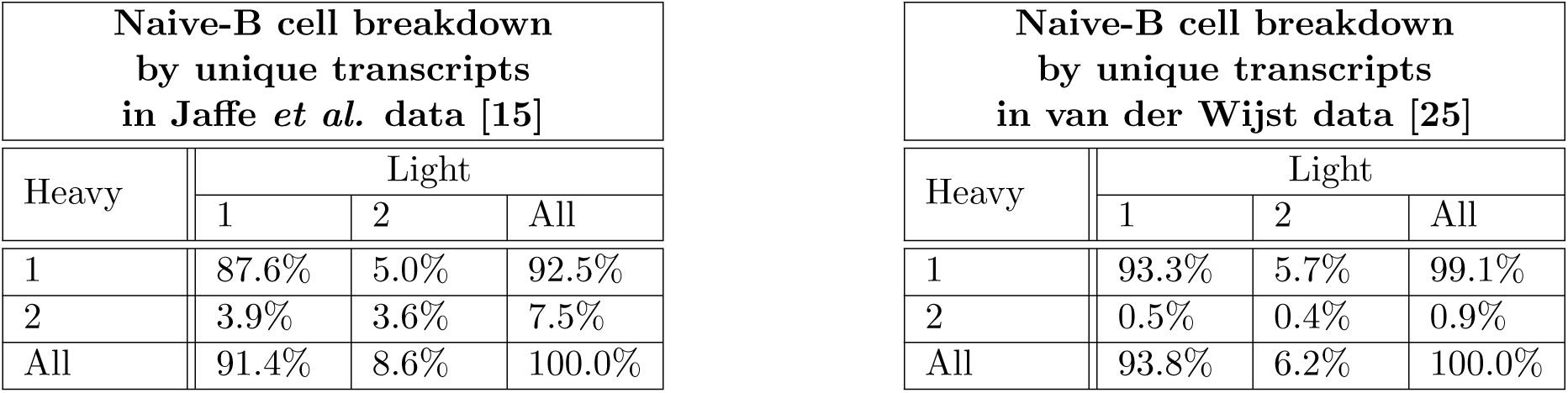
Observing allelic inclusion in single-cell BCR repertoires. (Left) Breakdown of barcodes in the Jaffe *et al.* dataset [15] by how many productive heavy and light chain transcripts they are associated with, in naive-B cells only. This dataset has not undergone computational multiplet filtering. Most barcodes have one of each, which is the expected standard allelic exclusion behavior. Other barcodes may be a result of allelic inclusion or of multiplet errors. (Right) Breakdown of barcodes in the van der Wijst *et al.* dataset [25] by how many productive heavy and light chain transcripts they are associated with, in naive-B cells only. This smaller dataset has undergone computational multiplet filtering. We therefore expect that it more accurately reflects true allelic inclusion statistics.

| Naive-B cell breakdown<br>by unique transcripts<br>in Jaffe *et al.* data [15] |  |  |  |
| --- | --- | --- | --- |
| Heavy | Light |  |  |
|  | 1 | 2 | All |
| 1 | 87.6% | 5.0% | 92.5% |
| 2 | 3.9% | 3.6% | 7.5% |
| All | 91.4% | 8.6% | 100.0% |

Table 1: **Observing allelic inclusion in single-cell BCR repertoires.**
| Naive-B cell breakdown<br>by unique transcripts<br>in van der Wijst data [25] |  |  |  |
| --- | --- | --- | --- |
| Heavy | Light |  |  |
|  | 1 | 2 | All |
| 1 | 93.3% | 5.7% | 99.1% |
| 2 | 0.5% | 0.4% | 0.9% |
| All | 93.8% | 6.2% | 100.0% |

Previous literature has found that heavy chain allelic inclusion is very rare in naive-B cells, with a rate of less than 0.1% [2]. Since the filtered dataset has a low rate of barcodes with two heavy chains, we postulated that the vast majority of barcodes with two heavy chains in the unfiltered Jaffe *et al.* dataset are due to multiplet errors. To investigate this claim, we fit classification models to distinguish simulated multiplet errors from barcodes with two heavy chains (**Methods**). We found that these barcodes were nearly indistinguishable from simulated multiplets, supporting the hypothesis that this part of the data is almost all multiplets (Supplementary Table S1). We therefore restricted our remaining analysis to double-light barcodes. We show in the next section that double-light barcodes include a substantial portion with clearly abnormal BCR sequences.

### Learning to identify dysfunctional sequences in allelic inclusion

Although we have now collected double-light B cells, which each contain a dysfunctional BCR sequence, we do not know which of the two sequences in each cell is dysfunctional. We developed a new framework to train a machine learning model on double-light B cells without this information (Figure 2A, **Methods**). Briefly, we use a neural network model that takes in the heavy and light chain sequences and outputs a “functionality” score. We train the model such that, when given the two BCRs from a double-light B cell, the minimum score is lower than the minimum score from two standard B cells (which both always have a functional BCR). The scores from such a model should then be lower for dysfunctional BCRs, allowing us to identify sequence constraints. We trained our model on the Jaffe *et al.* data, which may contain multiplets. However, the presence of multiplets in the double-light B cells should not bias predictions, since multiplet BCRs are indistinguishable from those of standard B cells. Training on the van der Wijst *et al.* dataset is less effective because the dataset is much smaller, so we reserve it for evaluation.

**Figure 2:**
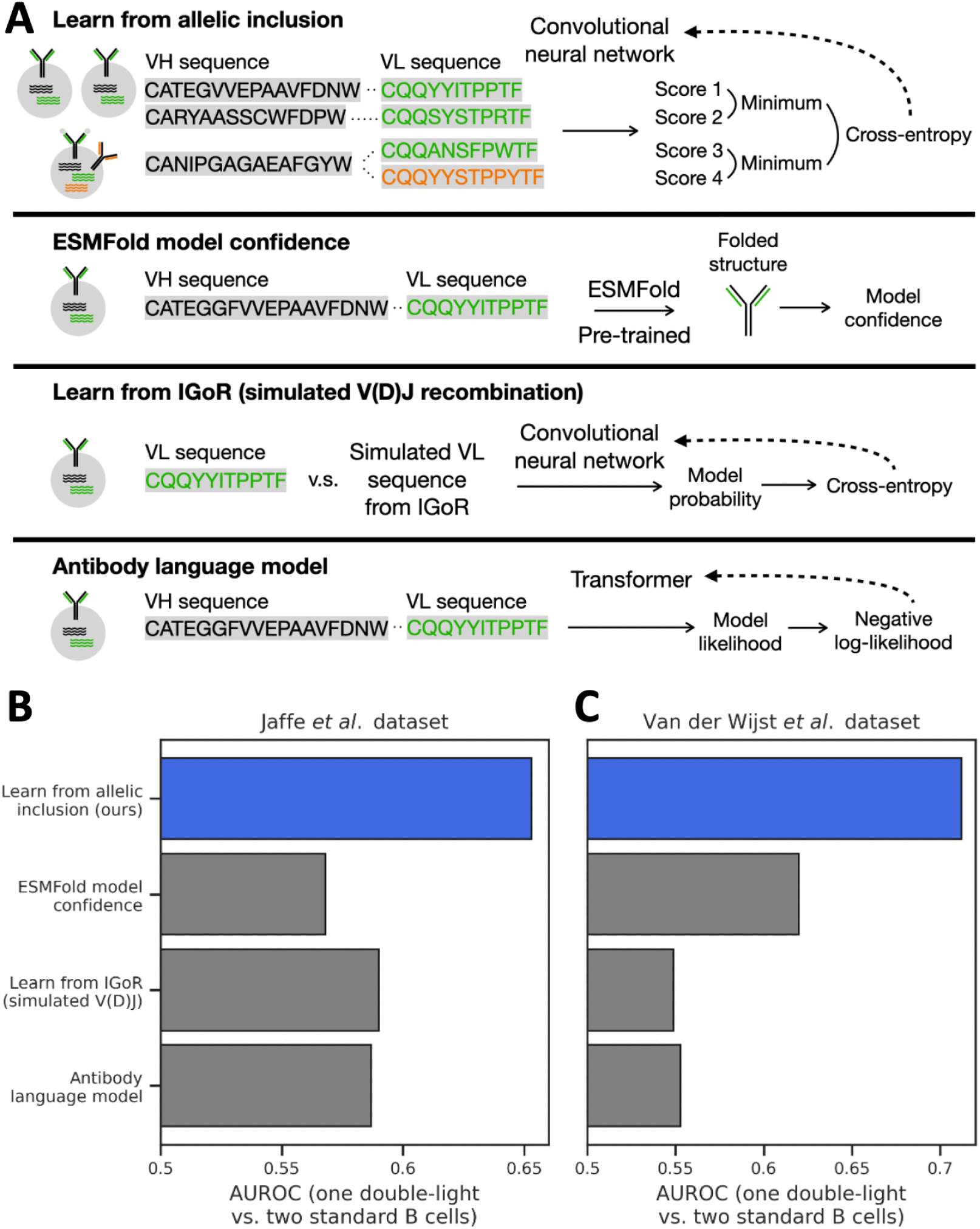
Learning to identify dysfunctional sequences in allelic inclusion. **(A)** Overview of the training procedure for our new model and baselines. We train our model to give a lower score to one of the two BCRs in double-light B cells, compared to the minimum score from two single-light B cells with functional BCRs. Dotted arrows point from a loss function to the model that is optimized to minimize that loss. ESMFold does not have a dotted arrow because it is pre-trained, and we do not modify it. See **Methods** for details. **(B-C)** We assess the ability of computational models to identify dysfunctional sequences in double-light barcodes. We find that our approach for learning from allelic inclusion improves accuracy over alternatives. Specifically, for each model, we compare the minimum score in double-light cells to the minimum score in pairs of standard cells. We assess on two datasets: (**B**) held-out donors in data from Jaffe *et al.* [15] and (**C**) a fully independent dataset from van der Wijst *et al.* [25].

We compared the accuracy of our new approach to that of three other approaches adapted suitably (Figure 2A). The adapted approaches are based on (1) ESMFold [26], a model that is similar to AlphaFold [27,28], (2) IGoR, a simulation of V(D)J recombination [23], and (3) antibody language models (**Methods**). We postulated that AlphaFold-based modeling could help distinguish structural differences between standard and dysfunctional BCRs, based on recent evidence that AlphaFold confidence metrics correlate with local energetic favorability [29, 30]. After initial experiments with AlphaFold, we switched to the much faster ESMFold for greater scalability after noticing that it achieved similar accuracy. We used IGoR to learn constraints on antibody sequence by simulating an unconstrained repertoire from V(D)J recombination and then contrasting this with naive-B-cell antibodies using a neural network classifier. We used antibody language models to score the functionality of antibody sequences based on the sequence likelihood according to the model. Our own novel approach learns directly from double-light barcodes in the Jaffe datasets. For a fair comparison with other approaches that do not directly use double-light barcodes as training data, we always assess our model on a separate donor not seen in training to ensure that the model is not specific to particular individuals. None of the models, including ours, is trained on any of the van der Wijst *et al.* data.

For each pair of heavy and light chain sequences, each machine learning method gives a score of how “good” the sequences are. These scores are obtained in different ways from different models, such as pLDDT from ESMFold, classification probability as a naive-B-cell sequence (as opposed to from simulated V(D)J recombination) from IGoR, sequence likelihood from language models, and classification score as functional from our method. To assess how well each method identifies dysfunctional BCR sequences, we compare scores from double-light B cells with scores from pairs of standard B cells. Specifically, we take the minimum score in each double-light cell and each pair of standard cells, then compute the AUROC between the two cell types. More accurate methods should have a higher AUROC, since true double-light B cells have a dysfunctional sequence but pairs of standard B cells do not. Note that this evaluation approach does not rely on knowing which sequence in each double-light cell is dysfunctional.

All four approaches tested have some predictive power to identify double-light cells, with our newly proposed approach being the most accurate by a significant margin on both datasets (Figure 2B-C). Among baseline models, ESMFold does as well as other approaches despite being trained on far fewer antibody sequences. The improved performance of our approach indicates that there is significant learnable signal in allelic inclusion data which is not captured by existing methods. The accuracy of all methods is limited by the presence of multiplets, especially in the Jaffe *et. al.* dataset.

Across both datasets, our model identifies over 5,000 double-light barcodes as possessing a clearly dysfunctional BCR sequence (less than 95% of scores in standard B-cells), which we interpret as likely to be true allelic inclusion instances. This is an order of magnitude more cells than existing analyses of allelic inclusion.

### Allelic inclusion model accuracy only depends on the light chain

We assessed which sequence features are most important for identifying dysfunctional sequences by taking away features from the model one by one (Supplementary Figure S1). This is known commonly as an “ablation” study in machine learning literature. Unexpectedly, we found that the model predominantly relies on the L chain sequence alone and does not use information from the paired H chain sequence, as removing the H chain does not decrease performance at identifying double-light cells. This indicates that double-light cells have a L chain which is generally dysfunctional, without a large dependence on the specific H chain of the cell. Note that this does not show that interactions between the H and L chains are unimportant; rather, it implies that *variability* in the H chain, such as at the CDRs, does not play a major role in L chain allelic inclusion. This is because our model can learn constraints imposed by non-varying parts of the heavy chain without explicitly taking the heavy chain as input. As a result of this finding, we hereafter use a version of the model that only takes the light chain as input. We return to model interpretation in a later subsection (**Interpreting learned constraints on antibody light-chain sequences**).

### Allelic inclusion model predicts antibody polyreactivity

We postulated that our model trained on allelic inclusion would correlate with light-chain sequence constraints. Concretely, we wondered whether it would correlate with specific properties associated with dysfunctionality, such as polyreactivity or surface expression. We next assess this idea on a few different datasets (Figure 3).

**Figure 3:**
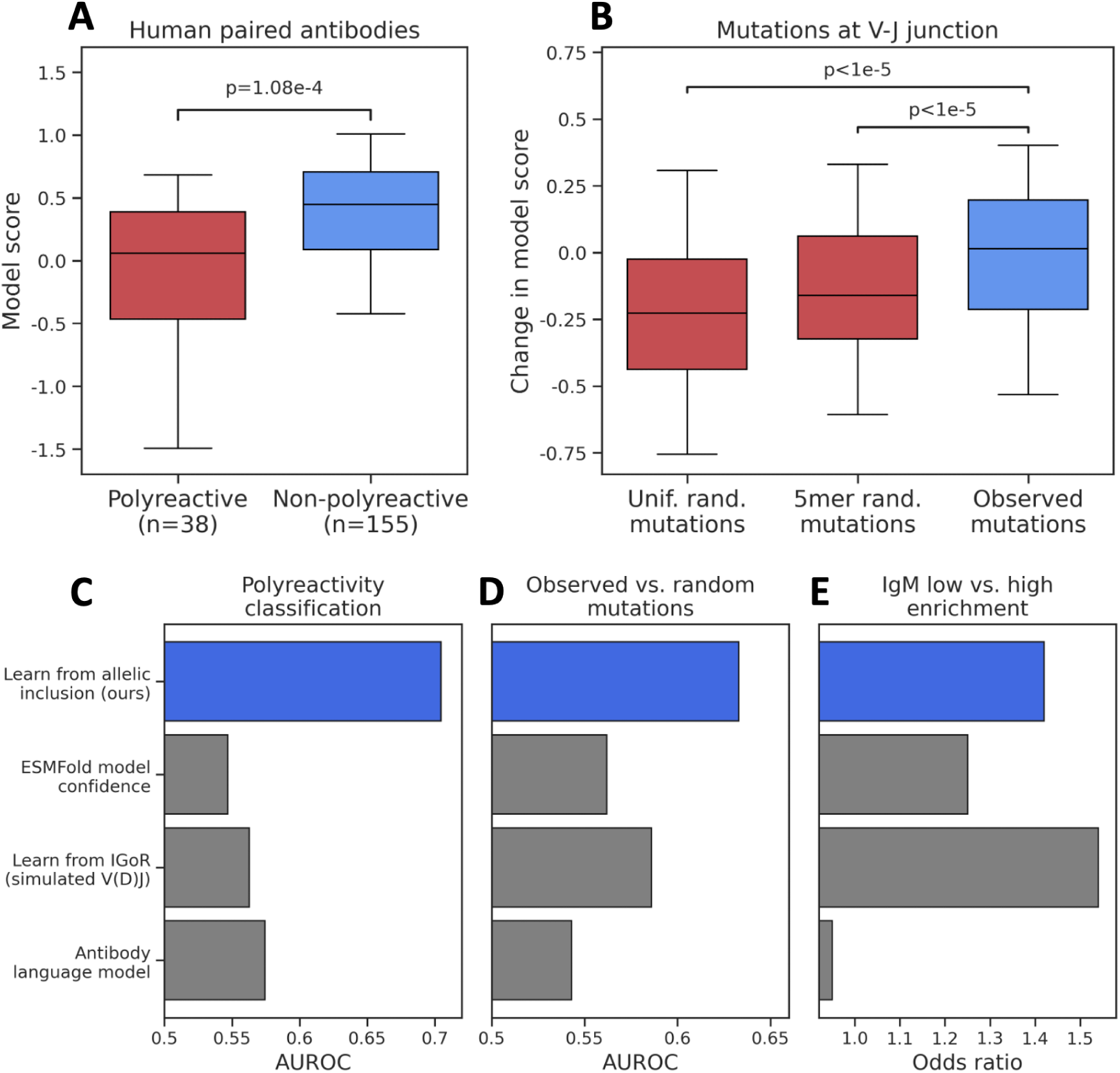
Allelic inclusion model predicts light-chain sequence constraints. (**A**) Our model assigns lower scores (associated with dysfunctionality) to polyreactive antibodies than those that are non-polyreactive. Data contains human paired antibodies from both before and after the autoreactivity checkpoint [4]. The model score is the log of the ratio of class 1 over class 0 for each sequence, where class 0 is double-light associated. P-value is calculated with a Mann-Whitney U test. (**B**) Mutations at the V-J junction that decrease the score from our model are depleted in affinity maturation. Observed mutations are collected from memory B cells. Control mutations are generated randomly at the same positions, either uniformly or based on a context-sensitive model of BCR mutation rates. We additionally restrict to mutations that occurred early in affinity maturation. P-values are calculated with a Mann-Whitney U test. We relax assumptions in Supplementary Figure S2. (**C-E**) We compared the allelic inclusion model with baseline antibody models and observed improved or comparable prediction accuracy. The allelic inclusion model discriminates polyreactive and non-polyreactive antibodies with much higher accuracy than other models. It discriminates observed versus random somatic mutations with higher accuracy (here, random is using 5-mer context). Enrichment of extreme scores in IgM low versus high is the second largest, behind the baseline model based on IGoR.

After light chain generation in small pre-B cells, most BCRs are autoreactive, whereas in mature naive-B cells, most BCRs are not autoreactive [2, 4]. We wondered whether our model, trained only on mature naive B-cells, would correlate with this reduction in autoreactivity observed in B-cell development. To test this idea, we used a dataset of human paired antibodies which were sequenced from both before and after the autoreactivity checkpoint and then assayed individually for polyreactivity [4] (**Methods**). After applying our allelic inclusion model to these data, we observed a clear correlation between antibody polyreactivity and model score, with polyreactive antibodies having lower scores (Figure 3A). This is particularly interesting since our model is only using the light chain, and adds to previous evidence indicating that choice of light chain is critical in determining whether an antibody is autoreactive [31]. The allelic inclusion model predicts polyreactivity much more accurately than baseline methods on this dataset (Figure 3C).

### Allelic inclusion model predicts mutation choice in affinity maturation

Beyond initial generation through V-J recombination, light chain sequences can be further modified through somatic hypermutation and selected for improved binding to a particular antigen (known as *affinity maturation*). However, during this process, constraints on expression and autoreactivity are still applied. We therefore wondered whether mutations that decrease scores from our model would be depleted in somatic mutations observed in memory B cells. A depletion would indicate that those mutations are associated with increased polyreactivity, reduced expression, or other negatively selected traits. We explored this idea using observed somatic mutations in memory B cells from the Jaffe *et al.* dataset [15].

Briefly, we reconstructed phylogenetic trees for each clone in the memory B cells and mapped light chain mutations to this tree. We then computed the change in model score from applying each mutation to the naive parent sequence of the light chain. As a control, we performed the same computation for uniformly random mutations at the same positions as well as mutations sampled from a model of BCR mutation rates conditioned on local sequence context [32]. We again only applied our models to held-out donors. We also note that our models were only trained on naive-B cells, but are tested in this application on memory B cells. For full details, see **Methods**.

We observed a difference in the change in score under our model between observed mutations and random mutations. Mutations that significantly decrease the model score are depleted in the observed mutation set. The effect is strongest if we restrict to mutations that occurred early in the reconstructed trees, as well as to mutations in the V-J junction (Figure 3B, **Methods**). The former assumption makes a difference because, as a clone accumulates many mutations, it moves out of distribution from the training data of our model (naive-B cells only). The latter assumption makes a difference for a similar reason; V-J recombination only generates complete amino acid diversity at the V-J junction. Therefore, many mutations at other sites are out of distribution from the training data of our model. We can remove the former assumption and observe a smaller effect on a much larger set of mutations (Supplementary Figure S2). However, for positions outside of the V-J junction, differences between distributions are very small. The allelic inclusion model is somewhat more accurate than baseline methods at discriminating observed and random mutations (Figure 3D).

### Low model scores are correlated with BCR surface expression

As a final application, we wondered whether our model scores would correlate with variation in BCR function across naive-B cells. Previous work has found that naive-B cells have large variation in surface IgM level (sIgM), and that cells with low sIgM have reduced antigen sensitivity and higher autoreactivity [33–35]. We wondered whether low scores from our model would be associated with the low sIgM phenotype, since such scores should be associated with impaired light chain function. Existing studies have not sequenced light chain repertoires stratified by this phenotype. To fill this gap, we collected new single-cell repertoires of paired BCR heavy and light chains, using naive-B cells stratified into IgM low and IgM high (**Methods**). We then applied our allelic inclusion models to all repertoires. We filtered out all double-light B cells to ensure these do not affect comparison of IgM low and IgM high compartments.

We observed an enrichment of very low model scores in the IgM low repertoires relative to IgM high. For example, at a threshold corresponding to the bottom 3% of IgM high sequences, scores below the threshold are enriched in IgM low by an odds ratio of 1.42 (*p* = 1.66 × 10*^−^*^3^, Fisher’s exact test). We do not observe significant enrichments for less extreme scores. This result indicates that the majority of dysfunctional light chains are successfully filtered out in B-cell development, but that a small fraction make it through and are subsequently associated with reduced surface expression in naive-B cells.

Compared to baseline antibody methods, the allelic inclusion model shows the second highest odds ratio on this data (Figure 3E). The model based on IGoR attains a higher odds ratio of 1.54 at a threshold corresponding to the bottom 3% of IgM high probabilities.

### Interpreting learned constraints on antibody light-chain sequences

The application datasets provide evidence that the allelic inclusion model learns constraints on antibody light chain sequences. We next looked to interpret the sequence features that the model associates with dysfunctionality (Figure 4). Such an analysis would not be possible without an accurate machine learning model, because we would not know which sequence in each true double-light cell is dysfunctional.

**Figure 4:**
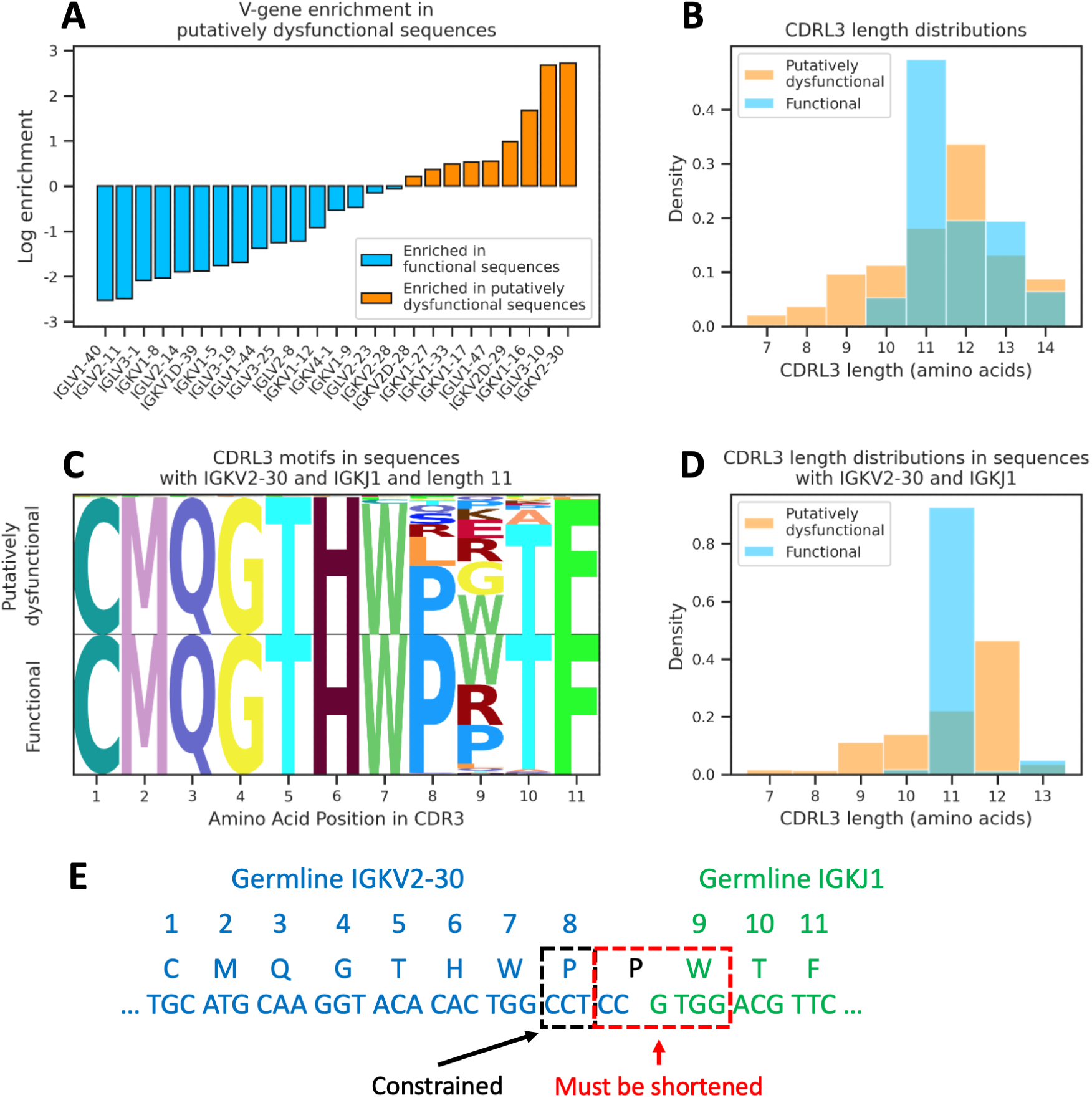
Interpreting learned constraints on light-chain sequences. Surprisingly, we found that model accuracy does not depend on the heavy chain (Supplementary Figure S1); we therefore focus on interpreting the light-chain sensitivity of our model. (**A**) Certain V-genes are highly enriched in dysfunctional light chains as predicted by our model, while others are highly enriched in functional light chains. Putatively dysfunctional light chains are those with a score less than 95% of scores in standard naive-B cells (**Methods**). (**B**) Short CDR3s and CDR3s of length 12 are enriched in dysfunctional light chains under our model (aggregated across all V- and J-genes). (**C-D**) We zoom in on IGKV2-30 and IGKJ1, a particular V- and J-gene with sufficient sample size in both the functional and putatively dysfunctional sets. We observe strong selection on CDRL3 length. Within the preferred length of 11, we observed significant differences at and near the V-J junction (position 9). Sample sizes in the functional and putatively dysfunctional sets are 747 and 491 across all lengths, and 634 and 72 within length 11. (**E**) The learned selection effects for this V- and J-gene pair can be interpreted by examining the germline sequences. To be functional, three nucleotides must be deleted during V-J recombination while preserving the surrounding nucleotides, which is a strict set of requirements.

We first calculated the enrichments of different light-chain V-genes in dysfunctional versus functional light-chains as predicted by our model (**Methods**). Enrichment is defined as the ratio of the frequencies in the two groups for each V-gene. We restricted to V-genes that have at least 1% usage in functional light-chains to ensure that we only analyzed those that have some chance of being functional. We observe that many V-genes are highly enriched in one group or the other (Figure 4A). We also observe strong selection on the length and amino acid composition of CDRL3 (Figure 4B, Supplementary Figure S3). In particular, short CDRL3s and CDRL3s of length 12 are enriched for dysfunctionality. However, the latter two analyses are confounded by varying V-gene usage between functional and putatively dysfunctional sequences.

To better understand what drives these enrichments, we examined predictions for a particular V- and J-gene (IGKV2-30 and IGKJ1) which are highly enriched for dysfunctionality but also have sufficient sample size in both groups (Figure 4C-D). We observe sharp selection on the length of CDRL3 (Figure 4D); nearly all functional sequences are length 11, while dysfunctional sequences are much more likely to be length 12. Within the preferred length of 11, we also find amino acid usage differences in the highly diverse V-J junction when contrasting functional and putatively dysfunctional sequences (Figure 4C). These learned patterns can be interpreted by examining the germline sequences for IGKV2-30 and IGKJ1 (Figure 4E). Our model predicts that CDRL3 should have length 11 to ensure functionality, but this requires three nucleotides to be deleted from the germline V- and J-genes during V-J recombination. In addition to this requirement, our model predicts that the nucleotides around these deletions must be somewhat conserved. These strict requirements help explain why this V-gene is enriched for dysfunctionality. Most V- and J-genes have insufficient sample size to enable such analysis. However, we verified that these general trends hold for a few more V- and J-gene pairs (Supplementary Figure S4).

### Early heavy chain repertoires contain sequences predicted to mispair

We have observed that selection on the light chain sequence is mostly independent of the paired heavy chain sequence, and that heavy chain selection does not play a major role in allelic inclusion in naive-B cells. The heavy chain is known to also be constrained by stability and autoreactivity, however, and we next sought to observe these constraints through other means. Allelic inclusion on the heavy chain has been observed at an earlier stage in pre-B cells [36]. However, due to the difficulty of sequencing the bone marrow, where pre-B cells reside, no large single-cell repertoire sequencing of pre-B heavy chains exists. This prevents us from conducting an analysis of heavy chains that is analagous to our light chain results.

Although large scale BCR repertoires from the bone marrow are not available in humans, previous work has collected such data in mice, with bulk instead of single-cell sequencing [6]. As an alternative approach to study heavy chain constraints, we used these data to compare naive-B-cell heavy chains with heavy chains from earlier development repertoires. During B-cell development, heavy chain rearrangement precedes light chain rearrangement, and a surrogate light chain (SLC) is expressed at the pre-B stage for screening pairing constraints on the heavy chain (Figure 5A). Between the pre-B and naive-B stages, the heavy chain is then screened for autoreactivity with a generated light chain. We looked to understand how SLC pairing and autoreactivity constrain the heavy chain by comparing heavy-chain repertoires at these three stages: VDJ recombination, pre-B cells, and naive-B cells. VDJ and pre-B repertoires have not passed all checkpoints and will therefore contain dysfunctional sequences. This approach does not require single-cell data, but is less precise because we cannot pick out the dysfunctional sequences via allelic inclusion. We used naive- and pre-B repertoires from C57BL6 mice [6] and simulated VDJ recombination repertoires from IGoR [23] (**Methods**).

**Figure 5:**
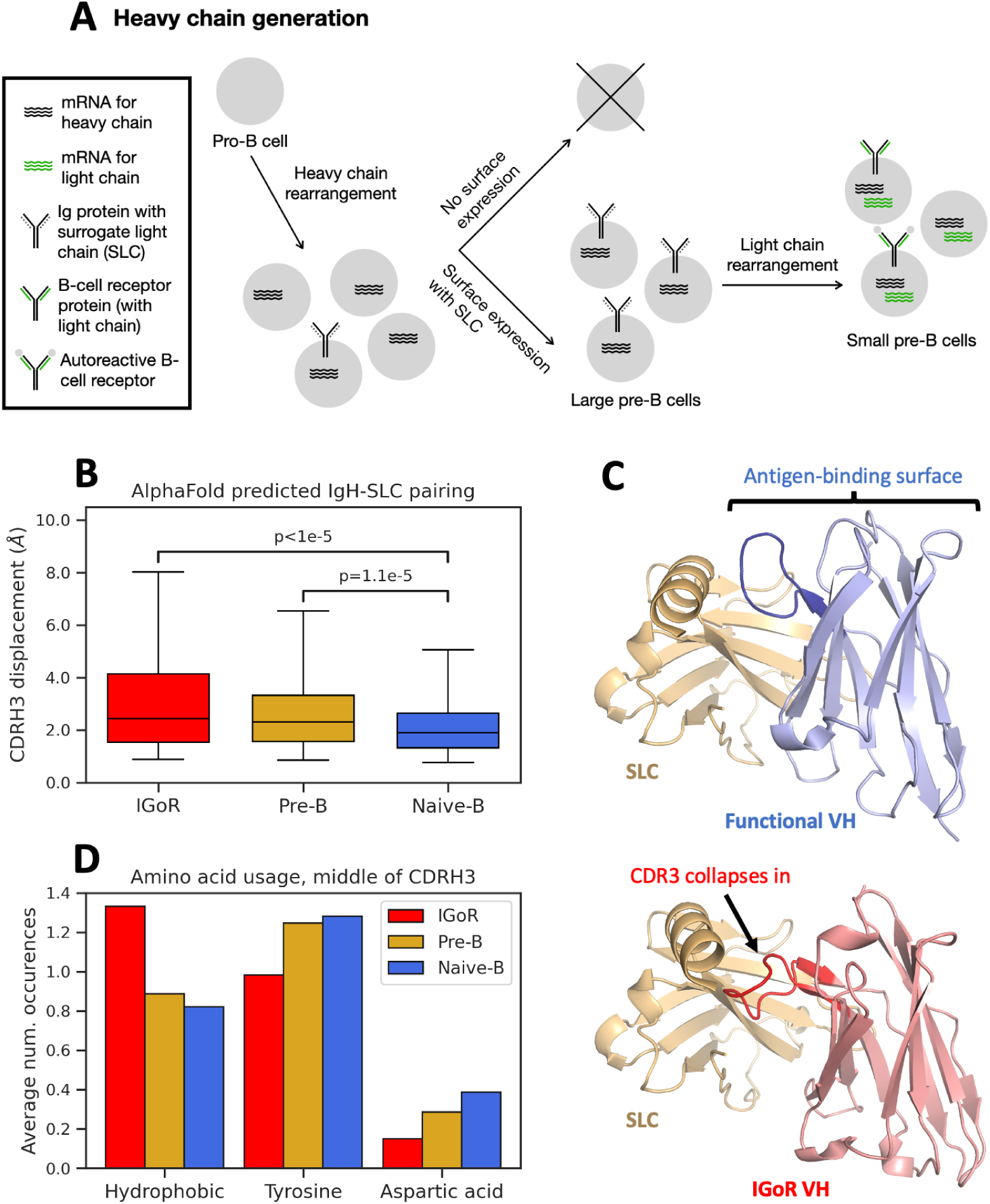
Surrogate light chain pairing constrains the heavy chain. (**A**) Overview of heavy chain generation. Heavy chains are generated in pro-B cells and tested for pairing with the surrogate light chain before progressing to pre-B cells. (**B**) Predicted heavy chain-SLC structures are abnormal much more frequently for IGoR heavy chains than naive-B heavy chains. Pre-B heavy chains are also predicted to form abnormal structures more frequently. Structures were predicted using AlphaFold, and structure abnormality is measured as the displacement of the heavy chain CDR3 from its typical location (**Methods**). P-values are from a Mann-Whitney U test. (**C**) We show example structures for correct (top) and incorrect (bottom) heavy chain-SLC structures. Functional heavy chains almost always have CDR3 in a particular location, whereas many IGoR heavy chains misplace CDR3. (**D**) Predicted structural differences appear to be driven by the amino acids in the D-gene region in the middle of CDR3 (**Methods**, Supplementary Figure S6). IGoR repertoires have higher usage of hydrophobic residues in this region. Naive repertoires use more tyrosine and aspartic acid.

Similar to our results on light-chain modeling, we found that AlphaFold-predicted structural interaction between the heavy chain and the SLC provides differential signals on the VDJ, pre-B, and naive-B repertoires. Simulated VDJ recombination sequences from IGoR are predicted by AlphaFold to form an incorrect structure in complex with the SLC much more frequently than naive- and pre-B sequences (Figure 5B). Specifically, the CDR3 in naive-B heavy chains is almost always placed in a specific location, but is often displaced in structures for IGoR-generated sequences (Figure 5C). Here, we calculate displacement as the distance between the the center of CDR3 and the typical location of the CDR3 center for naive-B heavy chains (**Methods**). Surprisingly, pre-B sequences also have higher displacement compared to naive-B sequences.

A higher predicted mispair rate among simulated VDJ recombination sequences reflects the fact that proper pairing with the SLC is enforced at the pre-B stage. However, it is unclear how to explain a higher rate of mispairing in the pre-B repertoire than the naive-B repertoire. We speculate that this is evidence of some pre-B cells displaying heavy chain allelic inclusion with one heavy chain not expressing on the surface, as has been previously observed [36]. We are unable to test this hypothesis at present because all sufficiently large pre-B-cell repertoires are from bulk sequencing.

When looking into what contributed to the observed differences in predicted structures, we found that IGoR heavy chain sequences are much more likely to have hydrophobic residues in the middle of CDR3, while pre-B and naive-B sequences replace these hydrophobic residues with residues such as tyrosine and aspartic acid (Figure 5D). Interestingly, these amino acid usages are mediated by the reading frame of the D-gene in CDR3. In many mouse D-genes, we observe one reading frame that contains a stop codon, one that is hydrophobic, and one that contains several tyrosines and aspartic acids (Supplementary Figure S5). IGoR heavy chains generally use all frames with significant probability, whereas pre- and naive-B heavy chains strongly prefer the last. These findings add to the evidence of previously speculated selection on the D-gene region of the heavy chain CDR3 sequence [37, 38].

### Machine learning models predict large restriction from heavy chain expression

We explored the possibility of training machine learning models to distinguish the three stages of mouse heavy chain repertoires (**Methods**). Concretely, we trained models to take a heavy chain sequence and predict whether it came from an IGoR, pre-B, or naive-B repertoire. Heavy-chain sequences that can confidently be predicted as non-naive are likely to fail development checkpoints. More specifically, sequences that are confidently predicted as coming from an IGoR repertoire are likely to not express, since all sequences in pre- and naive-B repertoires must have had adequate surface expression to progress. Sequences that express, in contrast, should be predicted as coming from pre- or naive-B cells with significant probability. Within sequences that express, those with a higher predicted probability of coming from a pre-B cell may be more autoreactive, since sequences are tested for autoreactivty between the pre- and naive-B stages.

We found that the vast majority of IGoR sequences are confidently predicted as being generated by IGoR, even if we only provide CDR3 to the model (Supplementary Figure S6). In contrast, pre- and naive-B heavy chain sequences can only be distinguished with modest accuracy. This indicates that cell surface expression constrains the heavy-chain repertoire more than autoreactivity. However, this claim depends on the reliability of simulated repertoires from IGoR.

### Machine learning models of mouse heavy chains correlate with polyreactivity

We next examined whether model predictions of pre-versus naive-B correlate with autoreactivity. We applied our model to a dataset of mouse antibodies that have been assayed for polyreactivity [39]. For each antibody in this dataset, we computed the predicted log odds of naive-to pre-B based on the heavy chain only (**Methods**). We observed a modest but statistically significant difference, with more polyreactive antibodies being associated with lower naive-B and higher pre-B probabilities (Supplementary Figure S7). No statistically significant correlation with polyreactivity is observed if we instead use a log odds to predicted IGoR probability. High polyreactivity will cause autoreactivity, so this result shows that the model can learn sequence determinants of heavy chain autoreactivity. However, we note that autoreactivity depends on the light chain, as we have seen in humans. Our mouse models only have access to the heavy chain in their training data and so their accuracy at identifying autoreactive sequences will be limited.

## Discussion

There has been broad interest in applying machine learning techniques to immune repertoire analysis, with applications in antibody engineering [40, 41] and clinical diagnostics [42, 43]. We have demonstrated that allelic inclusion in naive-B cells can be observed at sufficient scale to enable machine learning models of antibody sequence. Our results rely on recent advances in single-cell repertoire sequencing and depend on machine learning to identify the specific dysfunctional sequences in double-light B cells. Compared to baseline antibody modeling approaches, our new models showed improved correlation to properties like polyreactivity and mutation usage. This demonstrates the value of high-quality negative examples, compared to models trained only on positive examples [20–22, 26] or models trained with simulated negative examples [23, 24]. We also carried out an investigation into how similar forces constrain heavy chain sequences using bulk repertoires in mice. Beyond the new insights into B-cell development, our findings could be more generally integrated into machine learning approaches for antibody engineering and immune repertoire-based clinical diagnostics.

Our results suggest that constraints on heavy and light chain are largely applied independently. In particular, we found that the functionality of a light chain does not depend on which specific heavy chain it must pair with. Most constraints on the heavy chain, meanwhile, are enforced by the pre-B stage via the surrogate light chain, which does not depend on variability in the light chain. In addition, the specific constraints imposed on each chain are quite different. Heavy chain CDR3s showed evidence of strong selection against hydrophobicity in the D-gene region. Light chain CDR3s have particular lengths that are highly preferred and also preferred amino acids at the VJ junction. However, length of heavy chain CDR3 was not as tightly regulated, and light chain amino acid preferences do not show a clear effect due to hydrophobicity. Thus, despite their similarities, the differing structural roles of the heavy and light chain lead to significant differences in the selection forces affecting them.

Our allelic inclusion models learn a single overall functionality score that correlates with both polyreactivity and surface expression. However, the relationship between polyreactivity and surface expression is itself complex and unlikely to be fully captured by our models. For example, a CDR3 that is slightly more hydrophobic than usual may also be somewhat polyreactive. This could reduce surface expression by disrupting pairing, but also could increase polyreactivity even for molecules that make it to the surface. Experimental studies jointly measuring expression and binding properties of the same antibody sequences may help pull apart these factors. It is also interesting that antibody models have differing relative strengths on the application datasets we studied (Figure 3C). This may indicate that certain models correlate more with polyreactivity and others more with expression.

Among all the different sequence features tested, we observed that V and J genes alone are predictive of whether a light-chain sequence comes from an dysfunctional double-light cell (Supplementary Figure S1). It is possible that V and J gene usages may be biased by factors not related to biophysical functionality of a light chain sequence, such as the locality of the gene on the chromosome. Previous work has shown that downstream V and J genes are enriched in B cells that have used multiple rearrangements to generate a light chain [11, 12]. We observe this trend in *λ*-locus V gene usages, but do not see a significant effect in *κ*-locus V gene usages or J gene usages for either locus, and only saw slight shifts on *κ*- and *λ*-locus usage in double-light versus standard cells (Supplementary Figure S8). Even though V and J gene identities alone are predictive features, we found that other features regarding the biophysical determinants of antibody functionality, such as CDR3 length and CDR3 amino acid identities, further improve the model accuracy. Furthermore, the somatic mutation evaluation (Figure 3B) has an identical distribution of V and J genes between observed and control mutation sets, but we still observe significant predictive signal from our model.

Our study on allelic inclusion among light chains could be generalized in several directions. Allelic inclusion on the heavy chain has been observed in pre-B cells at small scale [36]. If larger single-cell sequencing repertoires of heavy chains from human pre-B cells become available, the same modeling approach could be applied there as well. Beyond B cells, allelic inclusion also exists in T cells and additionally interacts with MHC restriction [44–46].

## Methods

### Datasets

#### Single-cell human B-cell repertoires

From paired heavy and light chain sequencing data of 1.4 million B cells collected by Jaffe et al. [15], we train the allelic inclusion model on the subset of 448,650 cells marked as naive-B cells from flow cytometry and proceed with the same sequencing quality control filtering and V/J allele inference as in the original work. For Figure 4C-D and Supplementary Figure S4, we increased the sample size by additionally evaluating our model on sequences from cells in the pools labeled “naive + switched” and “naive + unswitched”. To restrict to naive sequences that have not undergone somatic hypermutation, we only kept sequences where the V-gene portion is unmutated from the germline. We additionally use the memory compartment of these data for analyzing somatic hypermutation.

We also use paired single cell BCR repertoires that were sequenced by van der Wijst *et al.* [25]. For these data, we recovered both gene expression vectors and BCR transcripts, and annotated cell types and doublets, using CellRanger as previously described [25]. We filtered out doublets and non-naive B cells, and then restricted to B cells with one heavy chain and at least one light chain. To apply models that were trained on the Jaffe *et al.* dataset, we further restricted to V-genes that were present in that data, which filtered out about 10% of remaining cells in the van der Wijst *et al.* dataset.

#### IGoR light-chain repertoire for human

IGoR (Inference and Generation Of Repertoires) [23] can infer V(D)J recombination simulation models from the statistics of unproductive BCR sequence reads in a repertoire, resting on the assumption that unproductive sequences do not interact with B-cell developmental checkpoints and thus reflect V(D)J recombination outcomes without additional selections. We generated IGoR light-chain repertoires for humans using the default IGoR light chain models for the human *κ* and *λ* loci. We then sampled from each locus at a proportion matching the *κ* - *λ* proportion in standard B cells in our single-cell human repertoire and filtered to only productive sequences. We did not have sufficient unproductive samples in the dataset to fit IGoR models specific to the individuals in our data.

#### Bulk C57BL/6 mouse B-cell repertoire

From heavy-chain only bulk sequencing data from mouse spleen and bone marrow collected by Greiff et al. [6], we focus on the pre-B-cell and naive-B-cell subsets from the antigen-naive C57BL/6 cohort of 5 mice. VDJ alignment and clonotype assembly were performed on each of the 10 repertoires by the MIXCR software [47] according to the IMGT [48] reference for the C57BL/6 strain. For downstream analysis of productive sequences, we filter for in-frame alignments with complete framework and CDR annotations from MIXCR and no stop codons.

#### IGoR repertoire generation for C57BL/6 mice

We generated IGoR heavy-chain repertoires for C57BL/6 mice. Our mouse dataset was of sufficient size to fit IGoR models for the specific mice in question. From each of the bulk C57BL/6 mouse naive-B-cell repertoires, we extract sequences that are annotated by MIXCR as having out-of-frame frameshift mutations in CDRH3 as data for an individual-specific IGoR model. We proceed to estimate the V(D)J recombination statistics with IGoR according to default hyper-parameters and the same IGMT reference as used in the earlier MIXCR annotation. Then, we sample 5 million simulated V(D)J recombination sequences from each IGoR model, followed by the same MIXCR annotation and filtering procedure as before.

#### ESMFold and AlphaFold structure generation

To generate predicted structures for human B-cell receptors, we use ESMFold version 1 [26], with chains connected by a linker of length 25. For mouse heavy chains together with the mouse surrogate light chain (SLC), we use AlphaFold Multimer [27] with templates but no multiple sequence alignments. In both cases, we predict structures with one copy of each variable domain, including all framework regions and CDRs. We use VPREB for the mouse surrogate light chain. Omitting the constant domains significantly increases speed and both models are able to infer the fold and positioning of variable domains without explicitly being given this context. For paired heavy and light chains, we superimpose all predicted structures to minimize the C*α* RMSD on the heavy chain variable region (excluding CDRH3). For heavy chains together with the mouse surrogate light chain (SLC), we superimpose to minimize the RMSD on the SLC, since the SLC sequence is identical in all structures. Initial experiments were run on ColabFold [49, 50]; subsequent structure predictions were run with local installations of AlphaFold and ESMFold. For human data, we generated 500 - 1500 predicted structures per application dataset. For mouse data, we generated 990 predicted structures, equally split between IGoR, pre-B, and naive-B heavy chain sequences.

#### Polyreactivity data for human paired antibodies

Polyreactivity data for human paired antibodies were obtained from Wardemann *et al.* [4]. This study sequenced antibodies from various stages of B-cell development and assayed those with a complete BCR in five ELISA-based binding tests (Insulin, ssDNA, dsDNA, LPS, nuclear antigen panel). We filtered to antibodies that were succesfully assayed against at least four of the five. We labeled as polyreactive those that showed binding in at least two and as non-polyreactive those that showed binding in none.

#### Expression-stratified BCR repertoires

B cells were isolated from 2 healthy PBMC donors using the EasySep™ Human B-cell Isolation Kit (Catalog #17954) from StemCell Technologies. Cells were then stained with APC anti-IgD-APC, anti-CD27-FITC, anti-CD19-PE, anti-IgM-APC-Cy7 antibodies and Zombie Violet Fixable Live/Dead stain, all from Biolegend, before IgM-high and IgM-low naive B cells (live, CD19+, CD27-, IgD+) were sorted on a BD FACSAria machine at the UCSF Parnassus Flow Cytometry Core. Following sorting, each sample was stained with TotalSeq C hashtags from Biolegend (Catalog #94661, 394663, 394667, 394671) according to Biolegend’s protocol. IgM-high and IgM-low thresholds were chosen to take the top and bottom 20% of naive B cells, respectively. Eighty thousand cells were then loaded in two Chromium Next GEM Single Cell 5’ v2 lanes from 10X Genomics and library prep was conducted with 10X’s BCR Amplification Kit (PN-1000253). Libraries were pooled and sequenced at a depth of 20,000 GEX reads per cell, 10,000 BCR reads per cell, and 5,000 Feature Barcode (hashtag) reads per cell at the UCSF Center for Advanced Technologies on a Novaseq X with 26×10×10×90 cycling parameters according to 10X’s recommendation. Data were processed with Cellranger v7 and were demultiplexed using the hashsolo library for hashtag information and freemuxlet for genetic information. We filtered out all doublets and cells with multiple light or multiple heavy chains, so that the expression analysis is not confounded by double-light B cells.

Although we sorted for CD27-, which is expected to remove memory B cells, we observed a large degree of clonal expansion in one of our two donors, especially in the IgM-low compartment. Concretely, for donor 2, 23% of IgM-low B cells and 9% of IgM-high B cells were in an expanded clone (at least three cells), compared to less than 0.5% in both compartments of donor 1. Analysis of gene expression vectors showed that clonally expanded cells had greatly elevated expression of certain HLA genes (HLA-A, HLA-DQA2, HLA-DRB5), as well as elevated expression of some conventional memory B-cell markers (CD80, CD86). CD27 was not differentially expressed, indicating that we may have isolated a population of CD27-memory B cells. Many other cells had similar expression patterns despite not being observed in expanded clones; such cells are likely to be antigen-experienced but not have had enough cells from the clone recovered in sequencing. To remove this antigen-experienced population, we filtered out all cells with expression of the aforementioned HLA genes greater than the 95th percentile of expression in donor 1; this filter removed over 99% of expanded clones. After this filtering, the reported odds ratios of low score enrichment are consistent between the two donors, and the reported statistic is from combining both together.

#### Mouse polyreactivity data

We applied our mouse heavy chain models to a dataset of mouse antibodies screened for polyreactivity [39]. These data include both heavy and light chains, but we only used the heavy chains since our models did not have access to light chain data. We defined high polyreactivity sequences as those that bound at least two out of seven antigens in the polyreactivity screen, and low polyreactivity sequences as those that bound none.

### Modeling and Analysis

#### Allelic inclusion model architecture and loss

For our analysis of allelic inclusion, we trained machine learning models that take a single antibody sequence (heavy and light chains) as input and predict whether it came from a standard or a double-light B cell. We used convolutional neural networks as the model architecture. The model uses two 2D convolutional layers on CDRH3 and CDRL3 which can learn interactions between the two chains. Specifically, both CDR3s are one-hot encoded, then zero-padded to length 32 and stacked to create a 2 × 32 × 20 input, with the one-hot dimension of length 20 being the channel dimension. The convolutional layers are followed by a single linear layer. The final output score therefore has a range of all real numbers. Earlier layers have ReLU activation. Before the linear layer, we concatenate the V- and J-genes for both chains with one-hot encoding. This is a small neural network; we could not use a more powerful model because of the limited number of double-light cells available.

To train our models, we used a non-standard loss function that makes full use of the single-cell resolution in our data. We expect that each double-light cell in our data contains at least one functional antibody sequence, because otherwise the B cell would not have survived to be sequenced. We also do not know beforehand which sequence is dysfunctional. We therefore constructed a loss function that only requires our model to predict one antibody sequence per double-light cell as dysfunctional, without knowing which one is bad *a priori*. At train time, we sample two standard B cells for each double-light cell. A double-light cell provides two antibody sequences (which have the same heavy chain but different light chains). The two standard B cells also provide two antibody sequences (which have different heavy and light chains). Our model architecture takes in a single sequence and outputs a score indicating whether the sequence comes from a standard B cell. We convert this real-valued score *x* to lie between 0 and 1 using the sigmoid transform *σ*(*x*) = (1 + exp(−*x*))*^−^*^1^. We then optimize the model to make the *minimum* output over the sequences from the double-light cell less than the minimum output over the sequences from two standard B cells. Mathematically, the loss function is a modified version of standard cross-entropy loss. Denote the heavy and light chains of the standard B cells as *h*_1_*, h*_2_*, l*_1_*, l*_2_. Denote the heavy and light chains of the double-light B cell as *h̄*, *l̄_1_*, *l̄*_2_. Denote the neural network as *f* and define *f_σ_*(*h, l*) := *σ*(*f* (*h, l*)). Then the contribution to loss function from these three cells is:

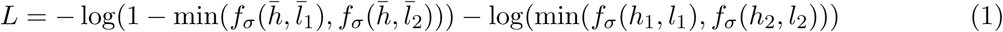

Due to the sigmoid, the argument of the logarithm is between 0 and 1 and the above equation can always be computed. At test time, we generally use the scores *f* (*h, l*) without applying a sigmoid, such that they are real-valued and not constrained to be between 0 and 1.

We assessed the benefit of this loss function by also training a model that simply classifies sequences as from a double-light versus a standard B cell. Our novel loss function improves accuracy both at identifying double-light cells with at least one dysfunctional sequence and at predicting double-light cells to have exactly one predicted dysfunctional sequence. Specifically, the fraction of double-light cells with at least one dysfunctional sequence at a 5% false positive rate improves from 27% to 31%, and the fraction of double-light cells with two sequences predicted as dysfunctional decreases from 1.5% to 1.1%. This behavior is more consistent with our knowledge of the B-cell biology. Intuitively, the conventional loss function incentives the model to predict both sequences in double-light cells as dysfunctional, which may skew the model towards relying on biases that are not directly connected to biophysical functionality.

We trained four separate models, each of which uses only three out of the four donors in our dataset for training. We then scored the entire dataset and, for each donor, used the model that was not trained on that donor. We tested other train-test splits, such as random splitting and splitting by sequence identity, and found that the held-out donor split was the most challenging (lowest performance for our model).

#### Double-heavy models and simulated multiplets

We additionally tested out training classifiers on barcodes with two heavy chains and either one or two light chains. For barocdes with two heavy chains and one light chain, the loss function is analagous to the above but with the roles of heavy and light chains swapped. For barcodes with two heavy and two light chains, there are several ways one could set up a predictive model. We tested (1) randomly pairing the two heavy and two light chains to get two paired instances (2) examining all four possible pairing instances (3) taking one heavy chain and both light chains and (4) taking one light chain and both heavy chains. The predictive performance was very similar in all four cases; we report results using approach (1). To generate Supplementary Table S1, we classify each population against simulated multiplets, constructed by combining two standard B cells, which each have one heavy and one light chain.

#### Allelic inclusion baselines and ablations

We compared our machine learning model to three baselines: ESMFold, an antibody language model, and a model trained on IGoR. For ESMFold, we first generated predicted interaction structure between the heavy and light chain variable domains (see **Datasets**). We then scored each antibody sequence using the average pLDDT of the light chain CDR3. We found that this feature was more predictive than several other features, including pLDDT of the heavy chain CDR3 and RMSD of the structure to a reference. The use of pLDDT was inspired by recent work by Roney and Ovchinnikov, which found that AlphaFold confidence metrics are correlated with the energetic favorability of a structure [29]. For Figure 2 and Figure 3, we estimated the discovery rates of ESM-Fold using 500 - 1500 predicted structures per application, because of the computational expense of applying ESMFold to the full datasets.

For antibody language modeling, we trained our own models on paired functional antibody sequences. While there are publicly available antibody language models [20–22], we wanted to train a model exclusively on the Jaffe *et al.* dataset to have a more controlled comparison of modeling approaches with other methods. Our training data included paired sequences from standard naive-B cells in the dataset, excluding double-light and double heavy cells. Training data are from different donors than the validation and test data. Each training example is a concatenated pair of a heavy chain sequence and a light chain sequence. Each sequence is the entire variable domain, including all framework regions and all CDRs. The models use an autoregressive transformer architecture and are trained using the fairseq [51] package. We tested various sizes (2-6 layers, embedding dimension 64-512) and reported results from the best among these hyperparameters. We then score sequences based on sequence log-likelihoods from the learned model.

For IGoR, we trained a model to predict enrichments of light chain sequences in standard B cells compared to IGoR light chain sequences (see **Datasets**). Specifically, the model is trained to classify light chain sequences from standard naive B cells (with one H and one L chain each) versus light chain sequences from IGoR (filtered to productive sequences only). This is a similar approach to the soNNia model, which builds on IGoR [24]. We used the same architecture as for our model trained on double-light cells, with the heavy chain portion of the input set to be constant. We did not incorporate heavy chains because no paired IGoR model is available and because we found for the allelic inclusion model that heavy-chain information is not useful. We also tested pretrained soNNia models for the light chain but found that these models did not support all the V-genes in our datasets. When restricting to V-genes that were supported, we found that pretrained soNNia models had very similar performance to our new baseline based on IGoR at classifying double-light cells.

Interestingly, we noticed that ESMFold pLDDT had some predictive signal on all datasets, but not always in the same direction. Concretely, for polyreactivity classification and IgM low versus high enrichment, the best sign is opposite of the best sign for allelic inclusion prediction. For ESMFold, we report results with the best sign on each application. Our new model and the model trained on IGoR have a consistent best sign on all four datasets. The antibody language model has a consistent best sign on three out of four and a statistically insignificant result in the other direction on the fourth, and so we do not flip its sign either.

We tested several ablations to our supervised model. To train a model that does not use the heavy chain, we used the exact same architecture but set all the heavy chain features to zero at train and test time. To train a model that only uses V-genes, J-genes, and CDR3 length from the light chain, we additionally set the CDRL3 sequence to be all alanines but keeping the length the same. To train a model that only uses V- and J-genes from the light chain, we trained a linear model on the one-hot encoded V- and J-genes.

#### Somatic hypermutation phylogenetic trees

We perform clonal family and germline inference with partis [52–54], which can exploit the paired heavy chain and light chain information provided in Jaffe *et al.* to form more accurate clonal clusters of BCR sequences. For each clonal family, we keep only productive sequences that do not have inferred insertions or deletions to avoid ambiguity of site positions. Using IQ-TREE, we then perform phylogenetic tree inference and ancestral sequence reconstruction on each clonal family [55], under a general time reversible substitution model and FreeRate site heterogeneity model, and taking the inferred naive sequence as outgroup. The heavy chain and light chain sequences are allowed to evolve at different rates under the *edge-linked-proportional* partition model [56].

To generate Figure 3B, we restricted the above data. First, we restricted to mutations that are mapped to edges coming from the naive parent. Mutations further down the tree will start to move out of distribution from the training data of our model, as BCR sequences move further from the space of possible outputs of V(D)J recombination. Second, we restricted to mutations at the V-J junction, defined as three positions back from the end of CDRL3. Other positions in the light chain do not have full amino acid diversity generated during V-J recombination, so many possible somatic mutations are out of distribution. We also tested relaxing these assumptions. If we do not filter to mutations coming from the naive parent, we observe smaller differences in the same direction between observed and random mutation model scores (Supplementary Figure S2). However, for positions outside of the V-J junction there are only very small effects.

Two control mutation sets were generated. For uniformly random mutations, we took the amino acid position of each observed mutation and uniformly sampled nucleotide positions and mutant nucleotides (with synonymous and stopgain variants set to zero probability). For local context dependent random mutations, we used 5-mer nucleotide contexts to adjust the probability of each mutation based on the HKL S5F model [32]. This model was inferred based on mutation patterns in productive light chains at sites that are 4-fold degenerate for the codon they belong to. These sites still experience somatic hypermutation but are not subject to selection because these are synonymous changes leaving the amino acid unchanged.

For each observed and control mutation, we computed the change in model score as the difference in score between the naive parent sequence and the naive sequence with the amino acid mutation applied.

#### Functional and putatively dysfunctional sequences

We visualized functional and putatively dysfunctional light chain sequences according to our model (Figure 4A-D, Supplementary Figure S4). Putatively dysfunctional sequences are sequences from a double-light cell with model score below 95% of scores in standard B cells. Functional sequences are sequences from a standard B cell with model score that is not below 95% of scores in standard B cells. For Figure 4A, log-enrichments are the natural log of the ratio of V-gene frequency in the two sets, restricting to V-genes with at least 1% usage in functional light chains.

#### Heavy-chain model architecture and loss

For heavy-chain modeling in mice, we trained machine learning models that take a single heavy chain as input and predict whether it came from IGoR, a pre-B cell, or a naive-B cell. The model uses four 1D convolutional layers on CDRH3, followed by two fully connected layers and a softmax. Before the fully connected layers, we concatenate the V- and J-genes with one-hot encoding. This model is significantly larger than our model for allelic inclusion, because the datasets are much larger. We trained our models with standard multi-class cross-entropy and provided a balanced distribution of the three classes at train time.

#### Analysis of heavy-chain AlphaFold predictions

We showed that IGoR heavy chains are predicted by AlphaFold to have a displaced CDR3 when interacting with the SLC more often than naive heavy chains (Figure 5B). To define a CDR3 displacement, we first calculated the mean location of CDR3 alpha-carbons for all structures, excluding the first three and last three positions. We then take the average of these CDR3 locations in naive heavy chain structures. The displacement for a particular CDR3 is then the distance between its mean location and the average in naive structures. We tested differences in the distribution of displacement using Mann-Whitney U tests.

#### Heavy chain amino acid and D-gene analysis

We analyzed amino acid and reading frame usage in the D-gene region of heavy chain CDR3s (Figure 5, Supplementary Figure S6). We calculated amino acid usages as the average number of occurrences of an amino acid in the middle five amino acids of CDRH3. We defined hydrophobic amino acids as I, V, L, F, C, M, and A based on the Kyte and Doolittle scale [57]. We extracted D-gene reading frames based on MIXCR annotations of the D-gene and its starting position.

## Acknowledgements

We would like to thank Antoine Koehl for helpful discussions. This research is supported in part by an NIH grant R35-GM134922. We also acknowledge support from Oracle Cloud credits and related resources provided by the Oracle for Research program.

## Supplementary Figures

**Figure S1:**
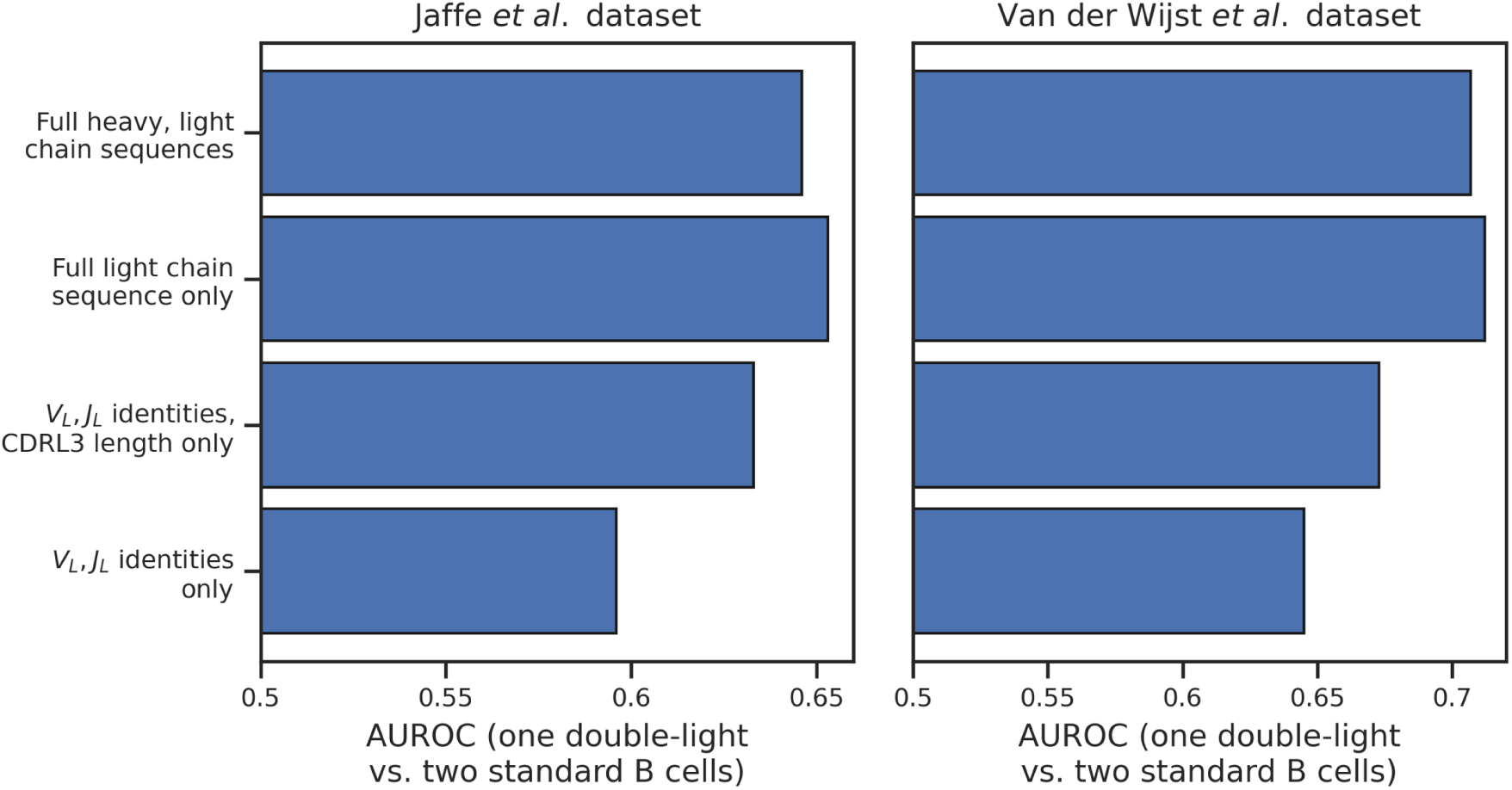
Feature ablations to allelic inclusion model. We tested removing different input features from our model. The accuracy of our model does not decrease when the heavy chain is removed. Within the light chain, removing CDR3 amino acids, and removing CDR3 length reduces performance further. A model that only sees *V_L_, J_L_* gene identities maintains significant accuracy, which is unsurprising since the full model predicts large differences in V-gene usages between normal and dysfunctional sequences.

**Figure S2:**
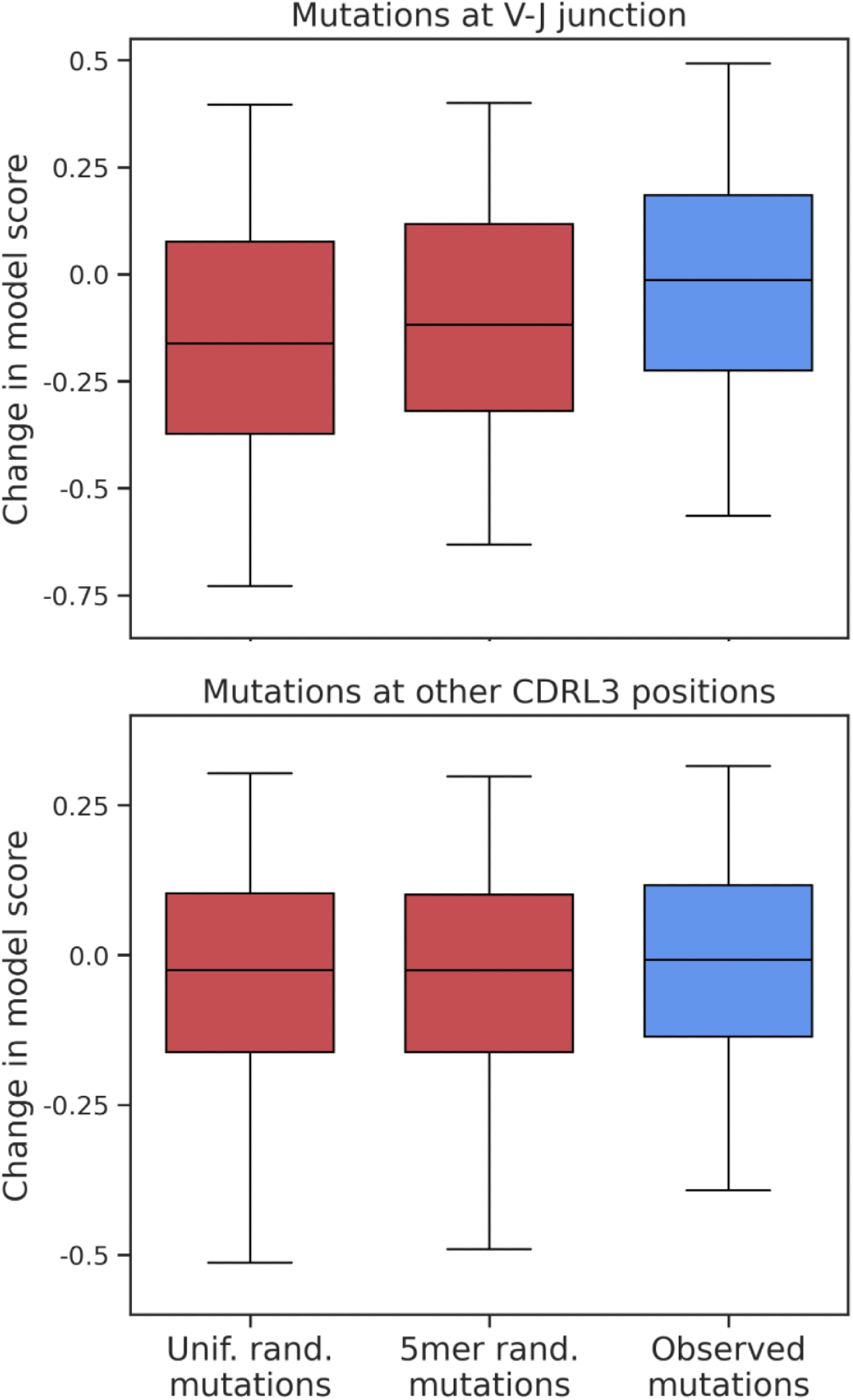
More general versions of somatic mutation bias. In Figure 3B, we showed a difference in model scores between observed and random somatic mutations, when restricting to mutations that occurred early in affinity maturation at the V-J junction. Here, we show the effect of removing these restrictions. (Top) We remove the first restriction, so that we keep mutations that occurred at any time in affinity maturation. A smaller effect in the same direction is observable over a much larger set of mutations (over 5000 versus around 500 in the original plot). (Bottom) We perform the same analysis on mutations at CDRL3 positions other than the V-J junction. Observed effects are much smaller.

**Figure S3:**
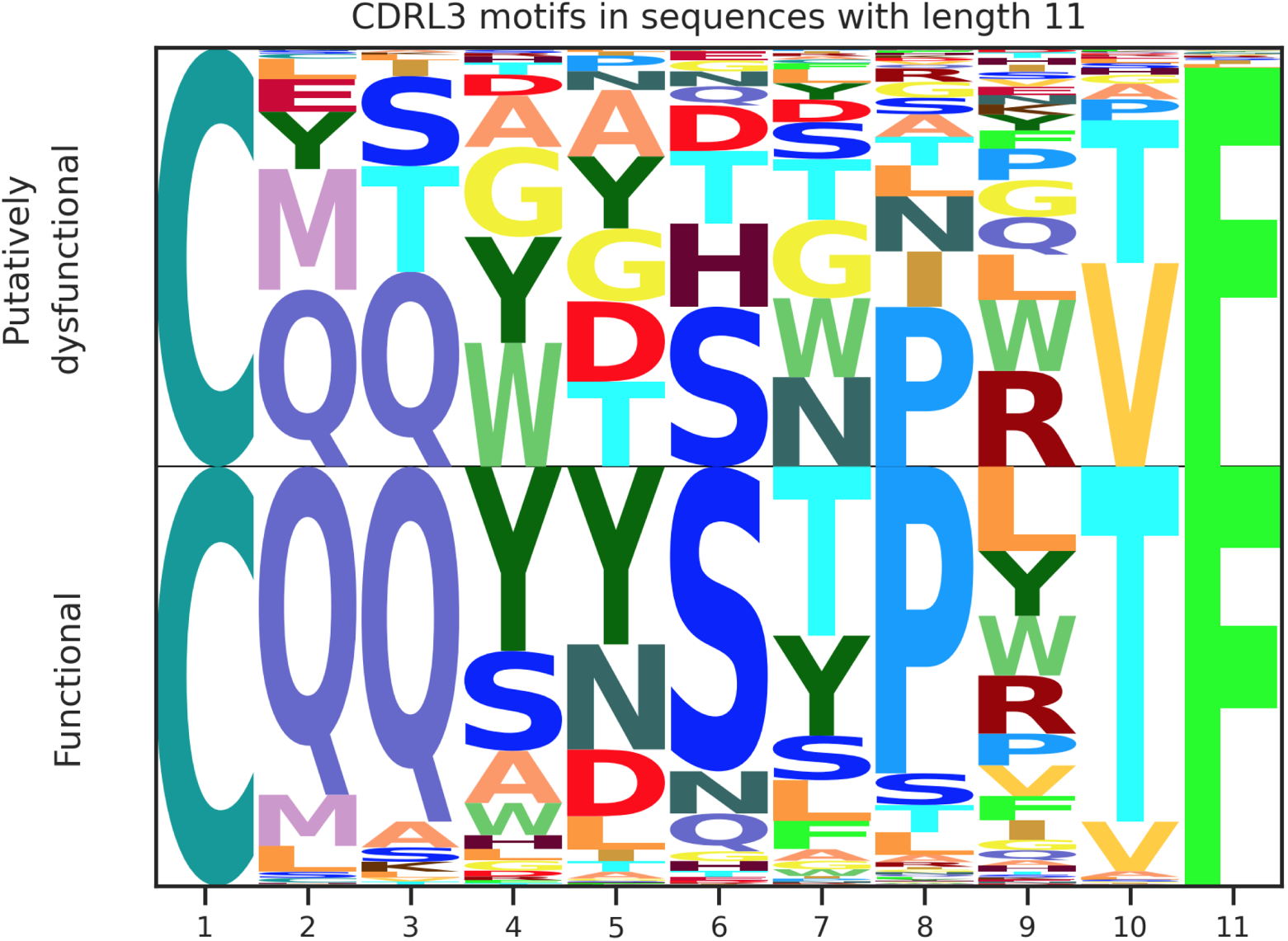
Amino acid usages in CDRL3. We compare sequence motifs for functional and putatively non-functional light chains in CDR3 conditioned on a particular length across all V- and J-genes. We observe clear biases, but also note that this analysis is confounded by varying V- and J-gene usages between the two sets.

**Figure S4:**
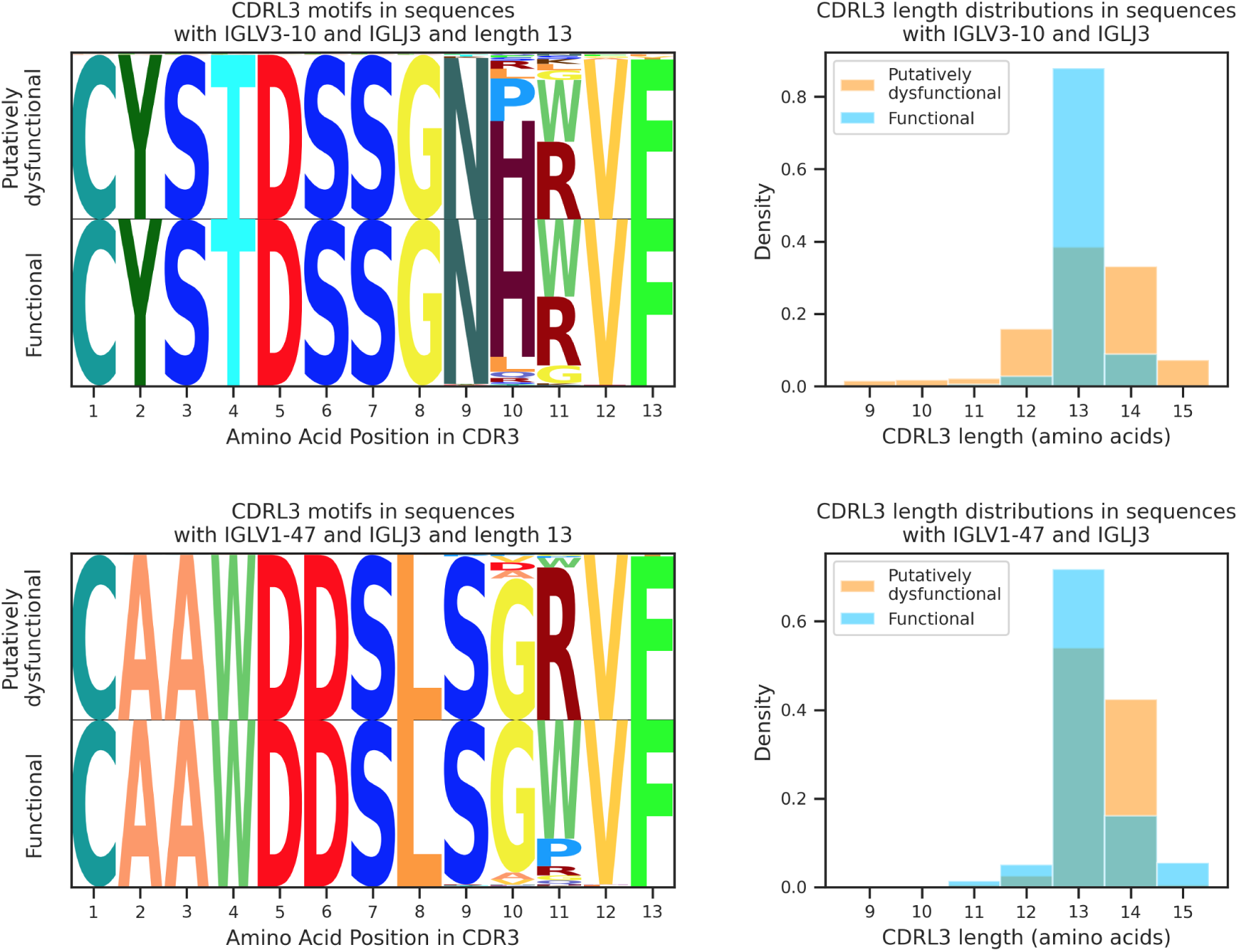
Functional and putatively non-functional sequences for more V- and J-gene pairs. We show length distributions and sequence motifs in the light chain CDR3 conditioned on two other V- and J-gene pairs, as in Figure 4C-D. We consistently observe strong length selection effects, although the preferred length varies by V-gene.

**Figure S5:**
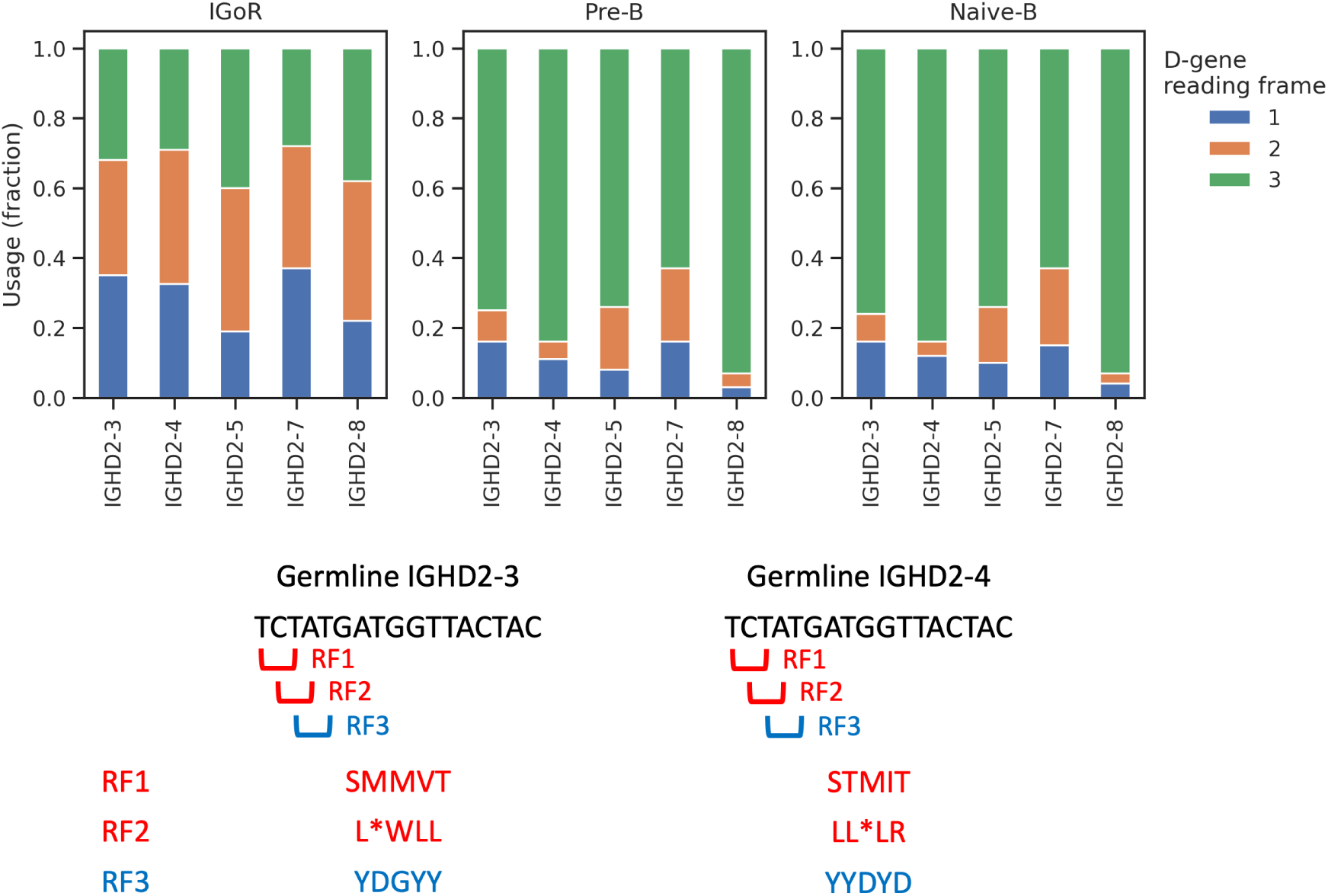
D-gene reading frames. (Top) In many mouse D-genes, there are large shifts in the reading frame usage from before (IGoR) to after (pre-B) selection for pairing with the SLC. The effects are largest for the IGHD2 family, which are used in about 40% of sequences in our dataset in all repertoire stages. (Bottom) We show the three D-gene forward reading frames for two D-genes (reverse frames are almost never used). Two of the three frames are negatively selected and have stop codons or high hydrophobicity. The third frame has several tyrosines and aspartic acids.

**Figure S6:**
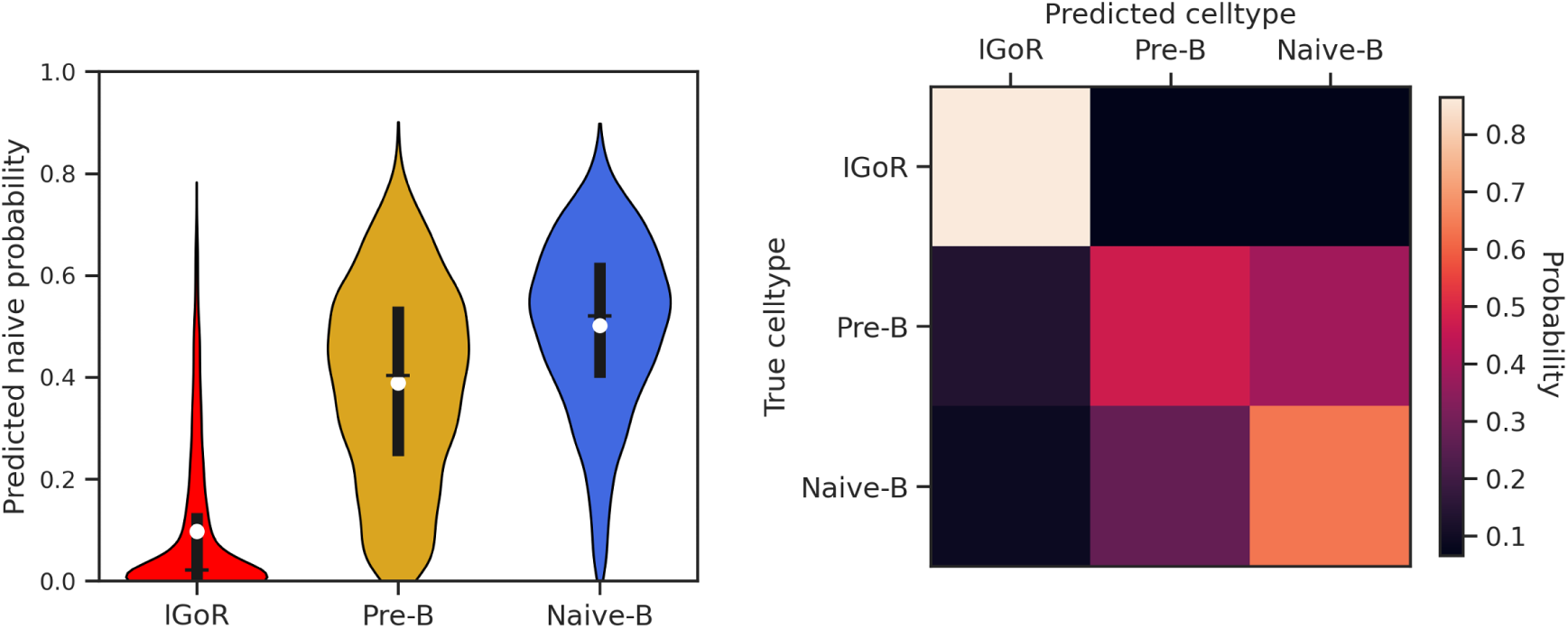
Machine learning on mouse heavy chains. We trained a machine learning model on antibody heavy chain sequences to predict selection stage. Nearly all IGoR sequences can be easily distinguished as abnormal, indicating that SLC pairing, which is enforced by the pre-B stage, significantly restricts the heavy chain repertoire. On the left, we show the distribution of predicted naive probability for all three classes, and on the right we show the 3-class confusion matrix.

**Figure S7:**
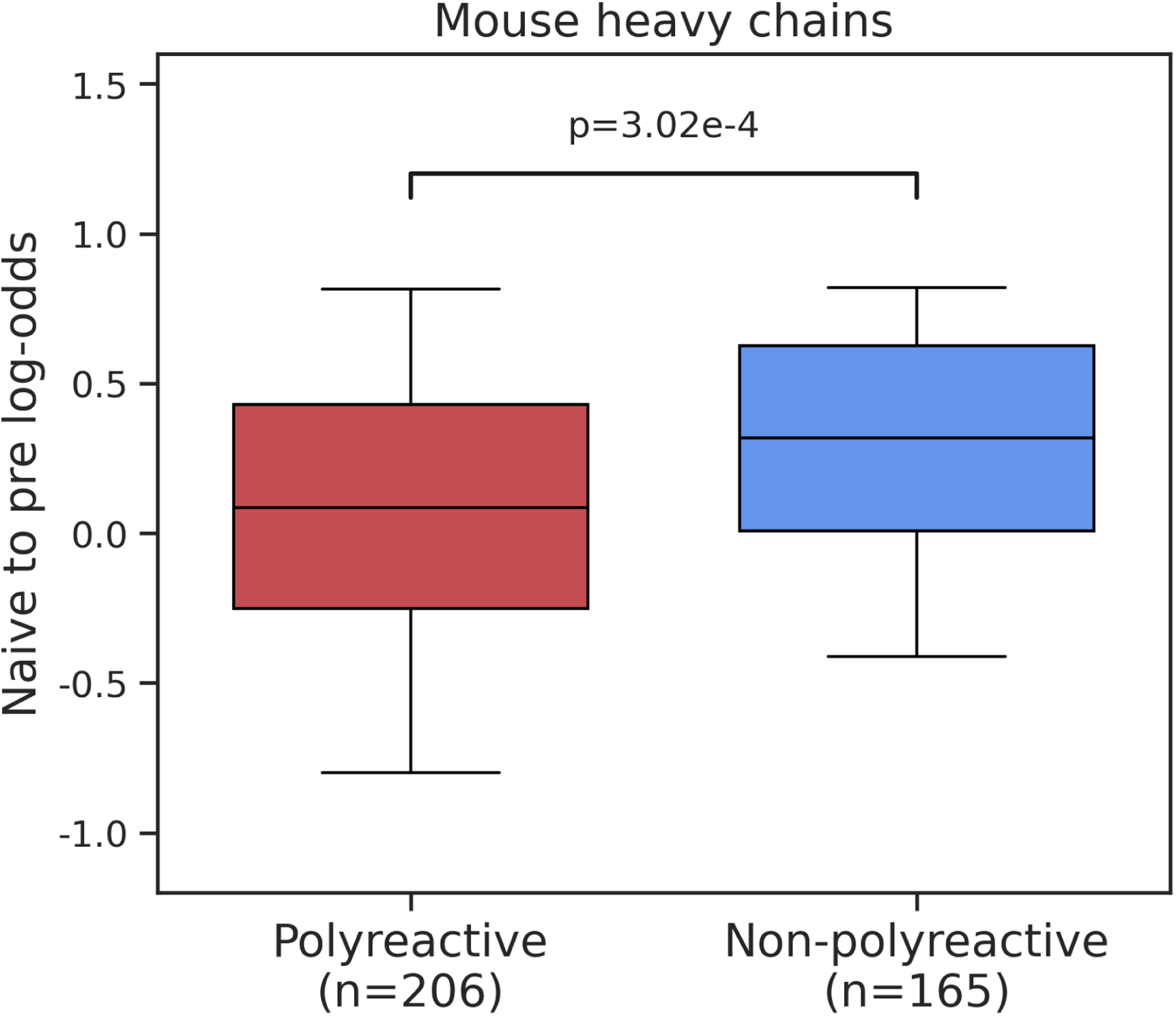
Mouse heavy chain polyreactivity. We applied our model of mouse heavy chain development to a dataset of mouse antibodies that were assayed for polyreactivity [39] (Methods). We found that polyreactive mouse antibodies were predicted to have lower naive to pre-B log odds from our model compared to non-polyreactive antibodies. P-value is computed with Mann-Whitney U test.

**Figure S8:**
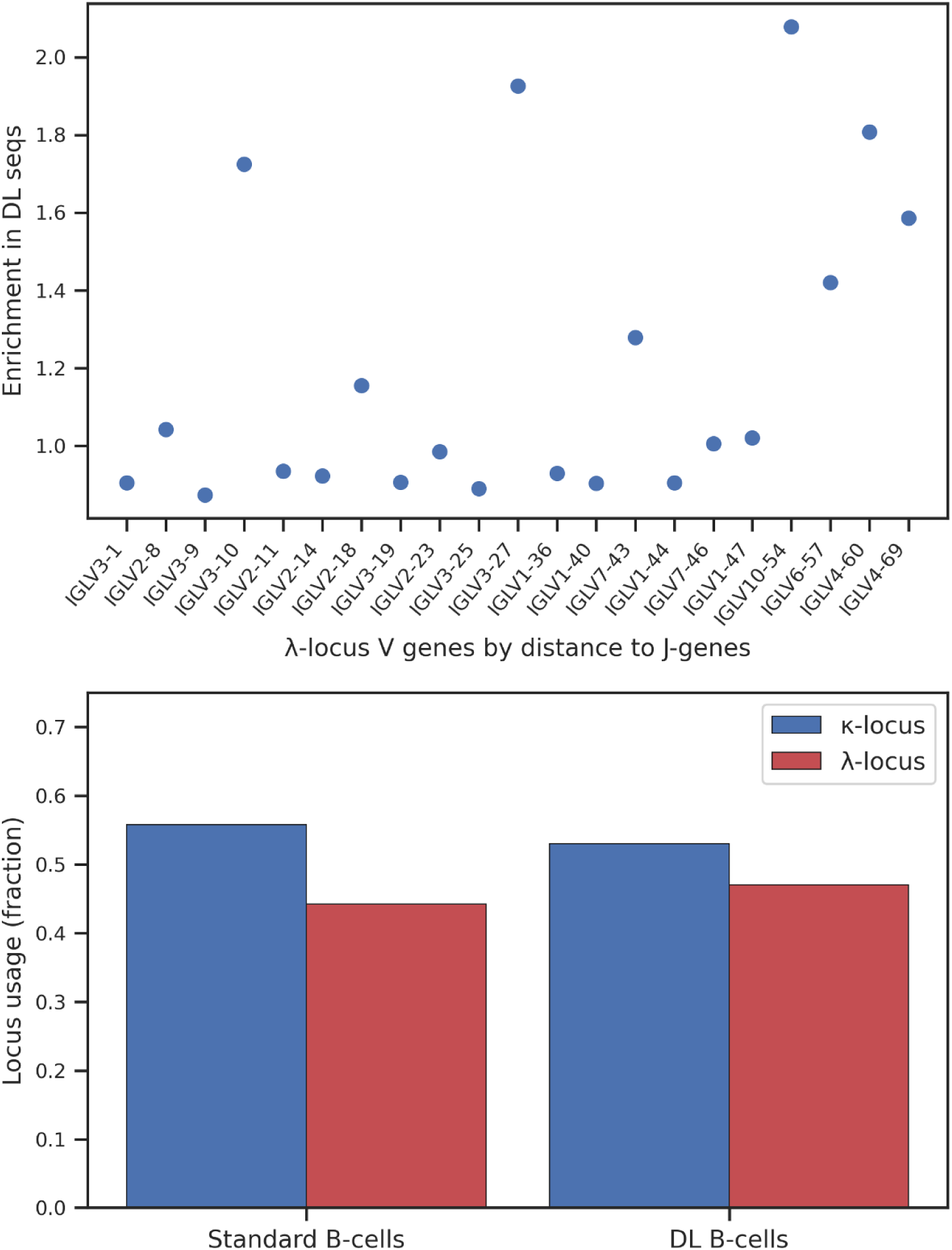
Gene usages. For both analyses, we used all DL barcodes, including multiplet errors. (**Top**) We calculated enrichment of light chain V- and J-genes in DL versus standard B-cell sequences as the ratio of the fraction of DL sequences over the ratio of standard sequences. We restricted to genes with at least 0.1% usage in standard sequences. We observed a statistically significant correlation between enrichment of *λ*-locus V-genes and their physical location in the genome. We did not observe a statistically significant effect for *κ*-locus V-genes or J-genes for either locus. This phenomenon has been previously reported and is a result of the mechansisms of receptor editing. (**Bottom**) *κ*-versus *λ*-locus usage only shows a minor shift between DL and standard B-cell sequences.

## Supplementary Tables

**Table S1:**
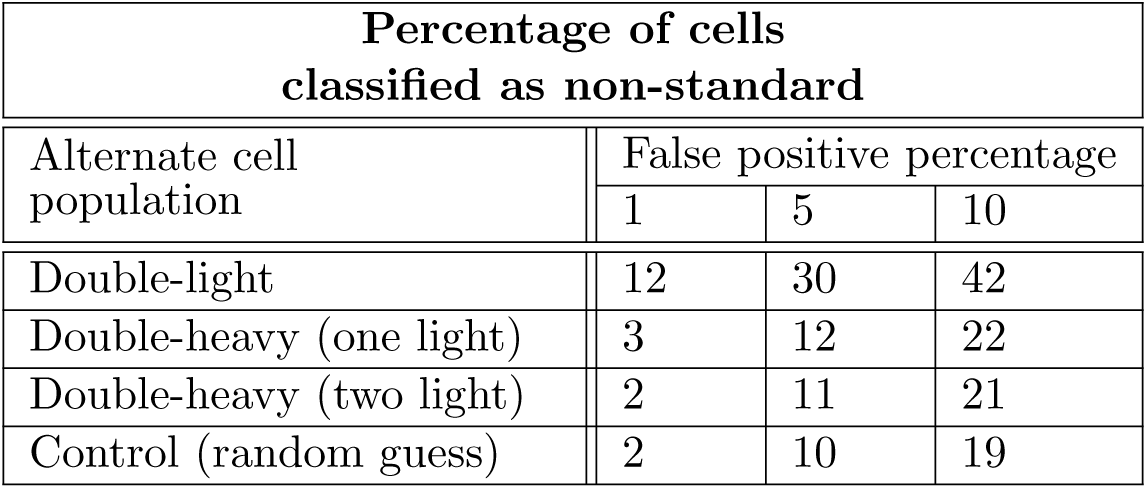
Classifying each multi-transcript population against simulated multiplets. We trained separate predictive models to distinguish each alternate population from simulated multiplets (pairs of standard B-cells) based on antibody sequence (**Methods**). The final row shows the percentages that would be predicted by random guessing as a control. On both types of double-heavy barcodes, the excess number of abnormal cells compared to control is very small, indicating that these populations are barely distinguishable to simulated multiplets. In contrast, double-light barcodes are predicted with an accuracy that implies a significant fraction of true allelically included cells in the population.

| Percentage of cells<br>classified as non-standard |  |  |  |
| --- | --- | --- | --- |
| Alternate cell<br>population | False positive percentage |  |  |
|  | 1 | 5 | 10 |
| Double-light | 12 | 30 | 42 |
| Double-heavy (one light) | 3 | 12 | 22 |
| Double-heavy (two light) | 2 | 11 | 21 |
| Control (random guess) | 2 | 10 | 19 |

## References

[1] William Hoffman, Fadi G Lakkis, and Geetha Chalasani. B cells, antibodies, and more. Clinical Journal of the American Society of Nephrology, 11(1):137–154, 2016.

[2] David Nemazee. Mechanisms of central tolerance for B cells. Nature Reviews Immunology, 17(5):281–294, 2017.

[3] Aaron L Nelson, Eugen Dhimolea, and Janice M Reichert. Development trends for human monoclonal antibody therapeutics. Nature Reviews Drug Discovery, 9(10):767–774, 2010.

[4] Hedda Wardemann, Sergey Yurasov, Anne Schaefer, James W Young, Eric Meffre, and Michel C Nussenzweig. Predominant autoantibody production by early human B cell precursors. Science, 301(5638):1374–1377, 2003.

[5] Ralf J Ludwig, Karen Vanhoorelbeke, Frank Leypoldt, Ziya Kaya, Katja Bieber, Sandra M McLachlan, Lars Komorowski, Jie Luo, Otavio Cabral-Marques, Christoph M Hammers, et al. Mechanisms of autoantibody-induced pathology. Frontiers in Immunology, 8:603, 2017.

[6] Victor Greiff, Ulrike Menzel, Enkelejda Miho, Cédric Weber, René Riedel, Skylar Cook, Atijeh Valai, Telma Lopes, Andreas Radbruch, Thomas H Winkler, et al. Systems analysis reveals high genetic and antigen-driven predetermination of antibody repertoires throughout B cell development. Cell Reports, 19(7):1467–1478, 2017.

[7] WR Strohl and LM Strohl. Development issues: antibody stability, developability, immunogenicity, and comparability. Therapeutic Antibody Engineering: Current and Future Advances Driving the Strongest Growth Area in the Pharma Industry (Strohl WR and Strohl LM eds) pp, 377:403, 2012.

[8] Paul J Carter and Arvind Rajpal. Designing antibodies as therapeutics. Cell, 185(15):2789– 2805, 2022.

[9] Edward P Harvey, Jung-Eun Shin, Meredith A Skiba, Genevieve R Nemeth, Joseph D Hurley, Alon Wellner, Ada Y Shaw, Victor G Miranda, Joseph K Min, Chang C Liu, et al. An in silico method to assess antibody fragment polyreactivity. Nature Communications, 13(1):1–15, 2022.

[10] Ameya Harmalkar, Roshan Rao, Yuxuan Richard Xie, Jonas Honer, Wibke Deisting, Jonas Anlahr, Anja Hoenig, Julia Czwikla, Eva Sienz-Widmann, Doris Rau, et al. Toward generalizable prediction of antibody thermostability using machine learning on sequence and structure features. MAbs, 15(1):2163584, 2023.

[11] Eline T Luning Prak, Marc Monestier, and Robert A Eisenberg. B cell receptor editing in tolerance and autoimmunity. Annals of the New York Academy of Sciences, 1217(1):96–121, 2011.

[12] Rafael Casellas, Qingzhao Zhang, Nai-Ying Zheng, Melissa D Mathias, Kenneth Smith, and Patrick C Wilson. Ig*κ* allelic inclusion is a consequence of receptor editing. The Journal of Experimental Medicine, 204(1):153–160, 2007.

[13] Valerie Kouskoff, Georges Lacaud, Kathryn Pape, Marc Retter, and David Nemazee. B cell receptor expression level determines the fate of developing B lymphocytes: receptor editing versus selection. Proceedings of the National Academy of Sciences, 97(13):7435–7439, 2000.

[14] Bryan Briney, Anne Inderbitzin, Collin Joyce, and Dennis R Burton. Commonality despite exceptional diversity in the baseline human antibody repertoire. Nature, 566(7744):393–397, 2019.

[15] David B Jaffe, Payam Shahi, Bruce A Adams, Ashley M Chrisman, Peter M Finnegan, Nandhini Raman, Ariel E Royall, FuNien Tsai, Thomas Vollbrecht, Daniel S Reyes, et al. Functional antibodies exhibit light chain coherence. Nature, 611(7935):352–357, 2022.

[16] Brandon J DeKosky, Oana I Lungu, Daechan Park, Erik L Johnson, Wissam Charab, Constantine Chrysostomou, Daisuke Kuroda, Andrew D Ellington, Gregory C Ippolito, Jeffrey J Gray, et al. Large-scale sequence and structural comparisons of human naive and antigen-experienced antibody repertoires. Proceedings of the National Academy of Sciences, 113(19):E2636–E2645, 2016.

[17] Brandon J DeKosky, Gregory C Ippolito, Ryan P Deschner, Jason J Lavinder, Yariv Wine, Brandon M Rawlings, Navin Varadarajan, Claudia Giesecke, Thomas Dörner, Sarah F Andrews, et al. High-throughput sequencing of the paired human immunoglobulin heavy and light chain repertoire. Nature Biotechnology, 31(2):166–169, 2013.

[18] Brandon J DeKosky, Takaaki Kojima, Alexa Rodin, Wissam Charab, Gregory C Ippolito, Andrew D Ellington, and George Georgiou. In-depth determination and analysis of the human paired heavy-and light-chain antibody repertoire. Nature Medicine, 21(1):86–91, 2015.

[19] Zhan Shi, Qingyang Zhang, Huige Yan, Ying Yang, Pingzhang Wang, Yixiao Zhang, Zhenling Deng, Meng Yu, Wenjing Zhou, Qianqian Wang, et al. More than one antibody of individual B cells revealed by single-cell immune profiling. Cell Discovery, 5(1):64, 2019.

[20] Jinwoo Leem, Laura S Mitchell, James HR Farmery, Justin Barton, and Jacob D Galson. Deciphering the language of antibodies using self-supervised learning. Patterns, 3(7):100513, 2022.

[21] Richard W Shuai, Jeffrey A Ruffolo, and Jeffrey J Gray. IgLM: Infilling language modeling for antibody sequence design. Cell Systems, 14(11):979–989, 2023.

[22] Tobias Hegelund Olsen, Iain H Moal, and Charlotte Deane. Addressing the antibody germline bias and its effect on language models for improved antibody design. bioRxiv, pages 2024–02, 2024.

[23] Quentin Marcou, Thierry Mora, and Aleksandra M Walczak. High-throughput immune repertoire analysis with IGoR. Nature Communications, 9(1):1–10, 2018.

[24] Giulio Isacchini, Aleksandra M Walczak, Thierry Mora, and Armita Nourmohammad. Deep generative selection models of T and B cell receptor repertoires with soNNia. Proceedings of the National Academy of Sciences, 118(14):e2023141118, 2021.

[25] Monique GP van der Wijst, Sara E Vazquez, George C Hartoularos, Paul Bastard, Tianna Grant, Raymund Bueno, David S Lee, John R Greenland, Yang Sun, Richard Perez, et al. Type i interferon autoantibodies are associated with systemic immune alterations in patients with COVID-19. Science Translational Medicine, 13(612):eabh2624, 2021.

[26] Zeming Lin, Halil Akin, Roshan Rao, Brian Hie, Zhongkai Zhu, Wenting Lu, Nikita Smetanin, Robert Verkuil, Ori Kabeli, Yaniv Shmueli, et al. Evolutionary-scale prediction of atomic-level protein structure with a language model. Science, 379(6637):1123–1130, 2023.

[27] Richard Evans, Michael O’Neill, Alexander Pritzel, Natasha Antropova, Andrew Senior, Tim Green, Augustin Žídek, Russ Bates, Sam Blackwell, Jason Yim, et al. Protein complex prediction with AlphaFold-Multimer. bioRxiv, pages 2021–10, 2022.

[28] John Jumper, Richard Evans, Alexander Pritzel, Tim Green, Michael Figurnov, Olaf Ronneberger, Kathryn Tunyasuvunakool, Russ Bates, Augustin Žídek, Anna Potapenko, et al. Highly accurate protein structure prediction with AlphaFold. Nature, 596(7873):583–589, 2021.

[29] James P. Roney and Sergey Ovchinnikov. State-of-the-art estimation of protein model accuracy using AlphaFold. Phys. Rev. Lett., 129:238101, Nov 2022.

[30] Philip Bradley. Structure-based prediction of T cell receptor: peptide-MHC interactions. Elife, 12:e82813, 2023.

[31] Hedda Wardemann, Johanna Hammersen, and Michel C Nussenzweig. Human autoantibody silencing by immunoglobulin light chains. The Journal of Experimental Medicine, 200(2):191, 2004.

[32] Ang Cui, Roberto Di Niro, Jason A Vander Heiden, Adrian W Briggs, Kris Adams, Tamara Gilbert, Kevin C O’Connor, Francois Vigneault, Mark J Shlomchik, and Steven H Kleinstein. A model of somatic hypermutation targeting in mice based on high-throughput Ig sequencing data. The Journal of Immunology, 197(9):3566–3574, 2016.

[33] T^am D Quách, Nataly Manjarrez-Orduño, Diana G Adlowitz, Lin Silver, Hongmei Yang, Chungwen Wei, Eric CB Milner, and Iñaki Sanz. Anergic responses characterize a large fraction of human autoreactive naive B cells expressing low levels of surface IgM. The Journal of Immunology, 186(8):4640–4648, 2011.

[34] J Andrew Duty, Peter Szodoray, Nai-Ying Zheng, Kristi A Koelsch, Qingzhao Zhang, Mike Swiatkowski, Melissa Mathias, Lori Garman, Christina Helms, Britt Nakken, et al. Functional anergy in a subpopulation of naive B cells from healthy humans that express autoreactive immunoglobulin receptors. The Journal of Experimental Medicine, 206(1):139, 2009.

[35] Diana Bautista, Camilo Vásquez, Paola Ayala-Ramcrez, Juan Téllez-Sosa, Ernestina Godoy- Lozano, Jesús Martínez-Barnetche, Manuel Franco, and Juana Angel. Differential expression of IgM and IgD discriminates two subpopulations of human circulating IgM+ IgD+ CD27+ B cells that differ phenotypically, functionally, and genetically. Frontiers in Immunology, 11:736, 2020.

[36] Edwin ten Boekel, Fritz Melchers, and Antonius G Rolink. Precursor B cells showing H chain allelic inclusion display allelic exclusion at the level of pre-B cell receptor surface expression. Immunity, 8(2):199–207, 1998.

[37] Mohamed Khass, Andre M Vale, Peter D Burrows, and Harry W Schroeder Jr. The sequences encoded by immunoglobulin diversity (DH) gene segments play key roles in controlling B-cell development, antigen-binding site diversity, and antibody production. Immunological Reviews, 284(1):106–119, 2018.

[38] Thomas H Winkler and Inga-Lill Mårtensson. The role of the pre-B cell receptor in B cell development, repertoire selection, and tolerance. Frontiers in Immunology, 9:2423, 2018.

[39] Christopher T Boughter, Marta T Borowska, Jenna J Guthmiller, Albert Bendelac, Patrick C Wilson, Benoit Roux, and Erin J Adams. Biochemical patterns of antibody polyreactivity revealed through a bioinformatics-based analysis of CDR loops. Elife, 9:e61393, 2020.

[40] Brian L Hie, Varun R Shanker, Duo Xu, Theodora UJ Bruun, Payton A Weidenbacher, Shaogeng Tang, Wesley Wu, John E Pak, and Peter S Kim. Efficient evolution of human antibodies from general protein language models. Nature Biotechnology, 2023.

[41] Raphael R Eguchi, Christian A Choe, Udit Parekh, Irene S Khalek, Michael D Ward, Neha Vithani, Gregory R Bowman, Joseph G Jardine, and Possu Huang. Deep generative design of epitope-specific binding proteins by latent conformation optimization. bioRxiv, pages 2022–12, 2022.

[42] Victor Greiff, Gur Yaari, and Lindsay G Cowell. Mining adaptive immune receptor repertoires for biological and clinical information using machine learning. Current Opinion in Systems Biology, 24:109–119, 2020.

[43] Maxim E Zaslavsky, Nikhil Ram-Mohan, Joel M Guthridge, Joan T Merrill, Jason D Goldman, Ji-Yeun Lee, Krishna M Roskin, Charlotte Cunningham-Rundles, M Anthony Moody, Barton F Haynes, et al. Disease diagnostics using machine learning of immune receptors. bioRxiv, 2022.

[44] Peter Borgulya, Hiroyuki Kishi, Yasushi Uematsu, and Harald von Boehmer. Exclusion and inclusion of *α* and *β* T cell receptor alleles. Cell, 69(3):529–537, 1992.

[45] Thomas Dupic, Quentin Marcou, Aleksandra M Walczak, and Thierry Mora. Genesis of the *αβ* T-cell receptor. PLoS Computational Biology, 15(3):e1006874, 2019.

[46] Giancarlo Croce, Sara Bobisse, Dana Léa Moreno, Julien Schmidt, Philippe Guillame, Alexandre Harari, and David Gfeller. Deep learning predictions of TCR-epitope interactions reveal epitope-specific chains in dual alpha T cells. Nature Communications, 15(1):3211, 2024.

[47] Dmitriy A Bolotin, Stanislav Poslavsky, Igor Mitrophanov, Mikhail Shugay, Ilgar Z Mamedov, Ekaterina V Putintseva, and Dmitriy M Chudakov. MiXCR: software for comprehensive adaptive immunity profiling. Nature Methods, 12(5):380–381, 2015.

[48] Marie-Paule Lefranc, Veronique Giudicelli, Chantal Ginestoux, Joumana Jabado-Michaloud, Geraldine Folch, Fatena Bellahcene, Yan Wu, Elodie Gemrot, Xavier Brochet, Jerome Lane, et al. IMGT®, the international immunogenetics information system®. Nucleic Acids Research, 37(suppl 1):D1006–D1012, 2009.

[49] Milot Mirdita, Konstantin Schütze, Yoshitaka Moriwaki, Lim Heo, Sergey Ovchinnikov, and Martin Steinegger. ColabFold: Making protein folding accessible to all. Nature Methods, 2022.

[50] Helen Berman, Kim Henrick, and Haruki Nakamura. Announcing the worldwide Protein Data Bank, 2003.

[51] Myle Ott, Sergey Edunov, Alexei Baevski, Angela Fan, Sam Gross, Nathan Ng, David Grangier, and Michael Auli. fairseq: A fast, extensible toolkit for sequence modeling. *arXiv preprint arXiv:1904.01038*, 2019.

[52] Duncan K Ralph and Frederick A Matsen, IV. Consistency of VDJ rearrangement and substitution parameters enables accurate B cell receptor sequence annotation. PLoS Computational Biology, 12(1):e1004409, 2016.

[53] Duncan K Ralph and Frederick A Matsen, IV. Likelihood-based inference of B cell clonal families. PLoS Computational Biology, 12(10):e1005086, 2016.

[54] Duncan K Ralph and Frederick A Matsen, IV. Per-sample immunoglobulin germline inference from B cell receptor deep sequencing data. PLoS Computational Biology, 15(7):e1007133, 2019.

[55] Bui Quang Minh, Heiko A Schmidt, Olga Chernomor, Dominik Schrempf, Michael D Woodhams, Arndt von Haeseler, and Robert Lanfear. IQ-TREE 2: New models and efficient methods for phylogenetic inference in the genomic era. Molecular Biology and Evolution, 37(5):1530–1534, 2020.

[56] Olga Chernomor, Arndt von Haeseler, and Bui Quang Minh. Terrace aware data structure for phylogenomic inference from supermatrices. Systematic Biology, 65(6):997–1008, 2016.

[57] Jack Kyte and Russell F Doolittle. A simple method for displaying the hydropathic character of a protein. Journal of Molecular Biology, 157(1):105–132, 1982.

